# Prokaryotic Single-Cell RNA Sequencing by *In Situ* Combinatorial Indexing

**DOI:** 10.1101/866244

**Authors:** Sydney B. Blattman, Wenyan Jiang, Panos Oikonomou, Saeed Tavazoie

**Author notes:** These authors contributed equally to this work (Sydney B. Blattman, Wenyan Jiang).

## Abstract

Despite longstanding appreciation of gene expression heterogeneity in isogenic bacterial populations, affordable and scalable technologies for studying single bacterial cells have been limited. While single-cell RNA sequencing (scRNA-seq) has revolutionized studies of transcriptional heterogeneity in diverse eukaryotic systems, application of scRNA-seq to prokaryotes has been hindered by their extremely low mRNA abundance, lack of mRNA polyadenylation, and thick cell walls. Here, we present Prokaryotic Expression-profiling by Tagging RNA *In Situ* and sequencing (PETRI-seq), a low-cost, high-throughput, prokaryotic scRNA-seq pipeline that overcomes these technical obstacles. PETRI-seq uses *in situ* combinatorial indexing to barcode transcripts from tens of thousands of cells in a single experiment. PETRI-seq captures single cell transcriptomes of Gram-negative and Gram-positive bacteria with high purity and low bias, with median capture rates >200 mRNAs/cell for exponentially growing *E. coli*. These characteristics enable robust discrimination of cell-states corresponding to different phases of growth. When applied to wild-type *S. aureus,* PETRI-seq revealed a rare sub-population of cells undergoing prophage induction. We anticipate broad utility of PETRI-seq in defining single-cell states and their dynamics in complex microbial communities.

## Main

Recent developments in high-throughput single-cell RNA sequencing (scRNA-seq) technology have enabled rapid characterization of cellular diversity within complex eukaryotic tissues^1–13^. Despite these advances, comparable tools to study the transcriptomes of individual bacterial cells^14–16^ have lagged behind significantly due to numerous technical challenges (Fig. S1). Current massively parallel eukaryotic scRNA-seq methods typically require custom microfluidics to co-encapsulate a single cell with a uniquely barcoded bead in a compartment, often a droplet^5, 6, 8^ or microwell^4, 7^. These approaches rely on two key properties of many eukaryotic cells, specifically that they are easily lysed with detergent to release their RNA and that their poly-adenylated mRNAs can be effectively captured by beads coated with poly(dT) primers. Adaptation of these approaches for bacteria is thwarted by the presence of thick prokaryotic cell wall^17^, which makes lysis challenging, and the lack of poly-adenylated mRNAs for effective capture.

Given these considerations, we identified *in situ* combinatorial indexing^18^ as an alternative basis upon which to develop a method for high-throughput prokaryotic scRNA-seq. Two conceptually similar eukaryotic methods, single-cell combinatorial indexing RNA sequencing (sci-RNA-seq)^11, 13^ and split-pool ligation-based transcriptome sequencing (SPLiT-seq)^12^, rely on cells themselves as compartments for barcoding, which abrogates the need for cell lysis in droplets or microwells. These methods are also amenable to reverse transcription (RT) with random hexamers instead of poly(dT) primers^12^. With just pipetting steps and no complex instruments, individual transcriptomes of hundreds of thousands of fixed cells are uniquely labeled by multiple rounds of splitting, barcoding, and pooling in microplates.

Here, we present Prokaryotic Expression-profiling by Tagging RNA *In situ* and sequencing (PETRI-seq), a high-throughput, affordable, and easy-to-perform scRNA-seq method capable of distinguishing the transcriptional states of tens of thousands of wild type Gram-positive (*S. aureus* USA300) and Gram-negative (*E. coli* MG1655) cells. PETRI-seq (Fig. 1) consists of three experimental components: cell preparation, split-pool barcoding, and library preparation, which are detailed in Figure S2 and Methods. Cells grown in liquid culture were briefly pelleted before fixation with 4% formaldehyde for 16 hours (Fig. S3A). We confirmed that centrifugation and fixation did not alter the bulk transcriptome (Fig. S3B,C). Cells were next resuspended in 50% ethanol, which has been used previously for prokaryotic *in situ* PCR as a storage solution^19^, though we have yet to test cellular and RNA integrity after long-term storage. Ethanol did not significantly change the cDNA yield from *in situ* RT (Fig. S3D). Lysozyme for *E. coli* (Fig. S3E), or lysostaphin for *S. aureus*, was subsequently added to permeabilize cells for *in situ* RT. Cells were next treated with DNase to remove background genomic DNA. We confirmed *in situ* DNase activity by qPCR (Fig. S3F) and verified DNase inactivation (Fig. S3G,H). DNase treatment did not significantly alter the bulk transcriptome (Fig. S3I) or RNA integrity (Fig. S3J). Before proceeding to RT, cells were imaged to confirm they were intact (Fig. S3K) and counted.

**Figure 1:**
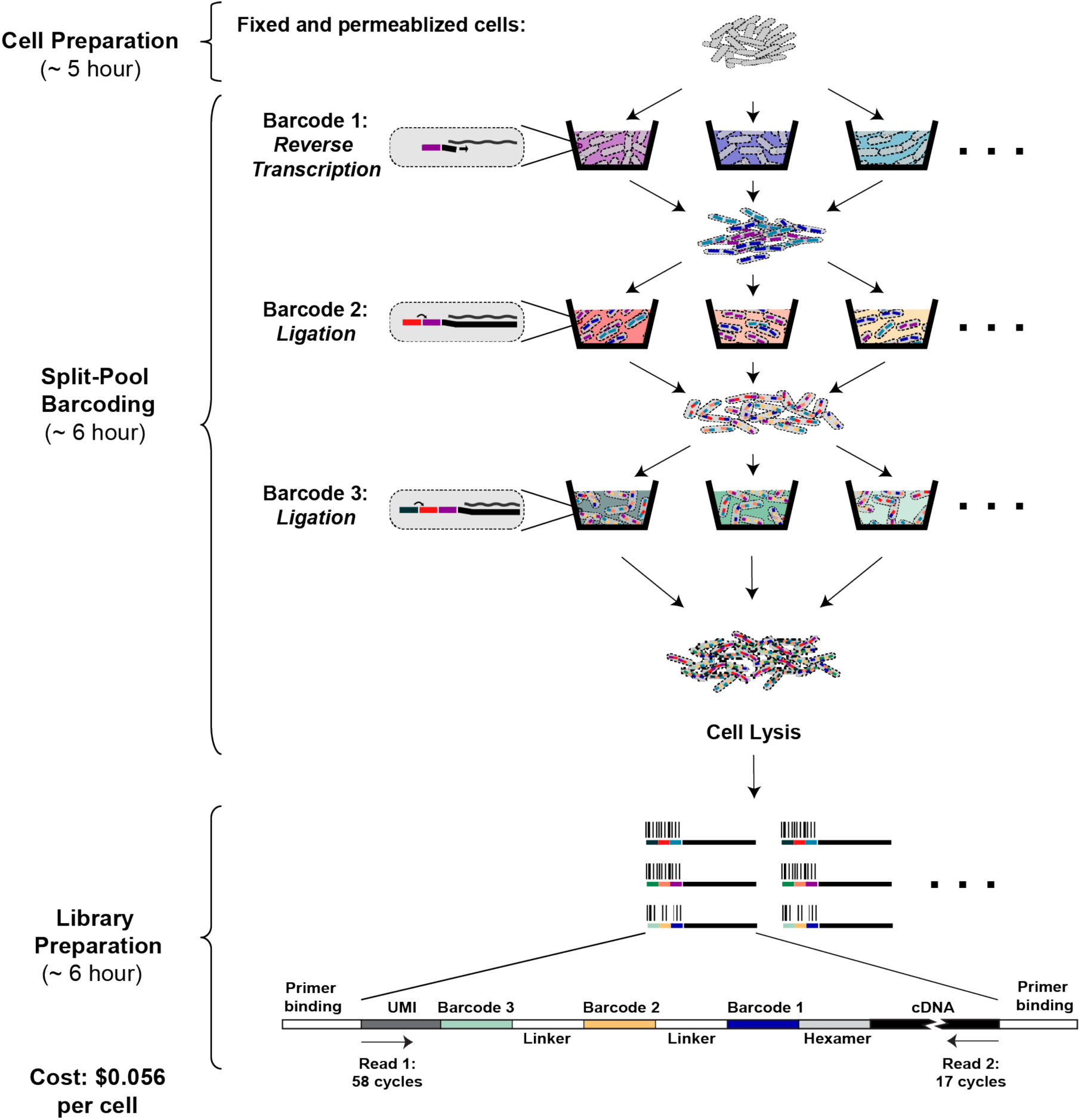
Overview of PETRI-seq. PETRI-seq includes three parts: cell preparation, split-pool barcoding, and library preparation. In cell preparation, cells are prepared for *in situ* reactions by fixation (formaldehyde) and permeabilization (lysozyme/lysostaphin). During split-pool barcoding, cells are split across 96-well plates three times for three rounds of barcoding by reverse transcription and two ligations. After barcoding, cells are lysed to release cDNA, which is subsequently prepared for paired-end Illumina sequencing. Each cDNA fragment in the library includes a unique molecular identifier (UMI) and 3 barcodes, which are all sequenced in Read 1. The UMI is a sequence of 7 degenerate nucleotides that can distinguish unique transcripts from PCR duplicates. The 3 barcodes comprise a barcode combination (BC), which allows reads to be grouped by their cell of origin. In Read 2, the cDNA is sequenced.

In the next stage, we performed split-pool barcoding (Fig. 1). Cells were distributed across a microplate for RT with random hexamers and different short DNA barcodes in each well. After RT, cells were pooled and redistributed across new microplates for two rounds of barcoding by ligation to the cDNA. We reduced the length of the overhang for each ligation relative to the eukaryotic protocol^12^, which made it possible to use only 75 cycles of sequencing instead of 150, thereby reducing sequencing cost by ∼50% (Table S1B). We demonstrated effective barcode ligation with this modification (Fig. S3L). After three rounds of barcoding, cells contained cDNA labeled with one of nearly one million possible three-barcode combinations (BCs). We counted the cells and lysed ∼10,000 cells for library preparation. The number of cells was chosen to ensure a low multiplet frequency, which is the percent of non-empty BCs containing more than one cell^20^. For a library of 10,000 cells, the expected multiplet frequency based on a Poisson distribution is 0.56%.

Finally, cDNA was prepared for Illumina sequencing (Fig. 1). We used AMPure XP beads to purify cDNA from cell lysates (Fig. S3M). AMPure purification is faster and significantly less costly (Table S1C) than primer biotinylation and streptavidin purification used previously in eukaryotic SPLiT-seq^12^. Next, to make double-stranded cDNA, we compared second-strand synthesis^21^ and limited-cycle PCR after template switching^2^. We found that the former had a significantly higher yield (Fig. S3N,O). We then performed tagmentation followed by PCR using the transposon-inserted sequence and the overhang upstream of the third barcode as primer sequences, thereby preventing amplification of any undigested genomic DNA. The libraries were sequenced and analyzed using the pipeline detailed in Figure S4 and Methods. We set a threshold based on total UMIs (unique molecular identifiers)^22^ per BC to distinguish cells from background (Fig. S4E,F).

To demonstrate the ability of PETRI-seq to capture transcriptomes of single cells, we performed a species-mixing experiment involving three populations of cells in exponential phase: GFP- and RFP-expressing *E. coli* and wild type *S. aureus* (Fig. 2A). From 14,975 sequenced BCs, we observed that BCs were highly species-specific with 99.8% clearly assigned to one species (Fig. 2B). We calculated an overall multiplet frequency of 1.5% after accounting for multiplets of the same species and non-equal representation of the two species^20^. Though this frequency exceeds the Poisson expectation of 0.85%, it is comparable to existing eukaryotic methods^8, 13^. Furthermore, within the *E. coli* population, BCs were highly strain-specific with 98.7% of plasmid-containing cells assigned to a single population (GFP or RFP) (Fig. 2C). While multiplet frequency is the probability of multiple cells travelling together during barcoding either by physical interaction or by chance, additional factors, such as barcoded free molecules released by occasional cell lysis, may compromise single-cell purity. This type of intercellular contamination has been described for eukaryotic scRNA-seq methods^23, 24^. To assess the contamination rate (the probability that a UMI in a single cell is derived from other cells) for PETRI-seq, we first excluded species-mixed multiplets and then found that BCs assigned as *E. coli* included a mean of 0.23% *S. aureus* UMIs (Fig. S5A, right), while BCs assigned as *S. aureus* also included a mean of 0.23% *E. coli* UMIs (Fig. S5B, right). After correcting for alignment ambiguities (Fig. S5E,F,I,J) and relative representation of the two species in the library, we calculated that 0.19-0.36% of UMIs in a PETRI-seq transcriptome were likely derived from other cells.

**Figure 2:**
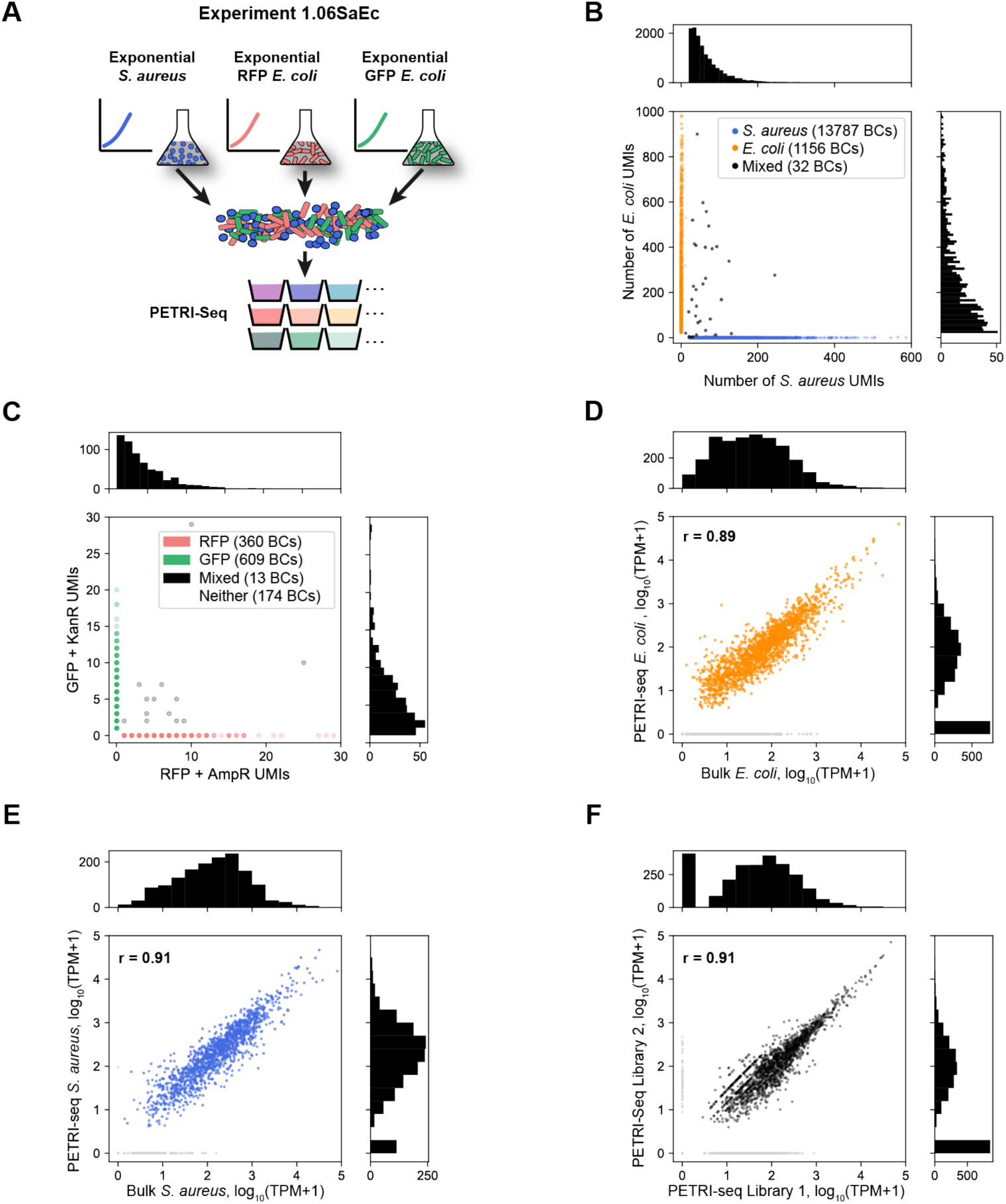
PETRI-Seq captures transcriptomes of single *E. coli* and *S. aureus* cells with high purity and low bias (**A**) Schematic of species-mixing experiment (1.06SaEc). Exponential *S. aureus* and *E. coli* cells were grown separately then mixed for PETRI-seq after cell preparation. *E. coli* cells included two populations containing either an RFP plasmid or a GFP plasmid, which were grown separately then combined for fixation. (**B**) Species mixing plot for *E. coli* and *S. aureus* based on total UMIs per BC, including rRNA. BCs were assigned to a single species if more than 90% of UMIs mapped to that species and fewer than 20 UMIs mapped to the other species. Histograms (*top, right*), show the number of *S. aureus* or *E. coli* cells (respectively) with the corresponding number of total UMIs. BCs with fewer than 20 total UMIs were omitted. The multiplet frequency is 1.5%. (**C**) Quantification of BC collisions within the *E. coli* population by plasmid mRNAs. Cells without plasmid genes (“Neither”) are omitted. BCs were assigned to a single cell type when greater than 90% of plasmid UMIs matched a single plasmid. Histograms (*top, right*) show the number of GFP BCs or RFP BCs, respectively, with the corresponding number of plasmid UMIs. (**D**) Correlation between mRNA abundances from PETRI-seq vs. a bulk library prepared from fixed *E. coli* cells. The Pearson correlation coefficient (r) was calculated for 1,873 out of 2,617 total operons, excluding those with zero counts in either library (grey points). If all operons are included, r = 0.78. (**E**) Correlation between mRNA abundances from PETRI-seq vs. a bulk library prepared from fixed *S. aureus* cells. Pearson’s r was calculated for 1,395 out of 1,510 total operons, excluding those with zero counts in either library (grey points). If all operons are included, r = 0.89. (**F**) Correlation between two biological replicate libraries of exponential GFP-expressing *E. coli* prepared by PETRI-seq. Pearson’s r was calculated for 1,714 out of 2,617 total operons, excluding those with zero counts in either library (grey points). If all operons are included, r = 0.78. For all correlations (E,F,G), PETRI-seq TPM was calculated from UMIs, and bulk TPM was calculated from reads.

Performing molecular reactions inside of cells raises the possibility that RNA capture could be biased by specific cellular contexts. Prior results in eukaryotic cells revealed a capture bias against rRNAs during *in situ* RT^12^, which is mildly recapitulated in our data (Fig. S6E,F). For exponential *E. coli*, 15% of sense PETRI-seq transcripts mapped to mRNA (Fig. S6E, pie chart), while only 5% of bulk sense transcripts mapped to mRNA (Fig. S6G, pie chart). Despite the capture bias against rRNA, we observed strong correlations between combined single-cell transcriptomes from PETRI-seq and bulk cDNA libraries prepared by standard RT for both *E. coli* and *S. aureus* (Fig. 2D,E). We also observed that reads mapped across the entire length of operons with minor bias against the 3’ end (Fig. S7A), which can be, at least, partially expected from our library preparation protocol (Fig. S7B). Our single-cell transcriptomes were reproducible, as shown by the strong correlation between the aggregated transcriptomes of GFP-expressing *E. coli* cells from two independent libraries (Fig. 2F).

Having confirmed that PETRI-seq captured transcriptomes of single cells with high purity and low bias, we next sought to determine the capacity of PETRI-seq to distinguish between cells in different growth states. In Experiment 1.10, we mixed *E. coli* cells in two well-characterized growth phases to create a population resembling naturally arising transcriptional heterogeneity. The mixed population consisted of GFP-expressing exponential and aTc-induced RFP-expressing stationary *E. coli* (Fig. 3A). We applied unsupervised dimensionality reduction (Principal Component Analysis—PCA^25^) to visualize the low-dimensional structure underlying the diversity of transcriptional states. For the PCA, we considered only cells containing at least 15 mRNAs to avoid spurious effects from cells with extremely low mRNA content. Without considering plasmid genes, we observed robust separation of two populations along principal component 1 (PC1). We then used the plasmid genes to classify these populations as RFP-containing stationary and GFP-containing exponential cells (Fig. 3B, bottom). We found that 98.5% of all plasmid-containing cells were on the expected side of an empirically chosen threshold line, and the threshold line predicted RFP cells with a 98.59% true positive rate (TPR) to the left of the line and GFP cells with a 98.5% TPR to the right. Of the 7,387 cells analyzed, 61% did not contain any plasmid transcripts, so their growth state was at first ambiguous (grey points in PCA).

**Figure 3:**
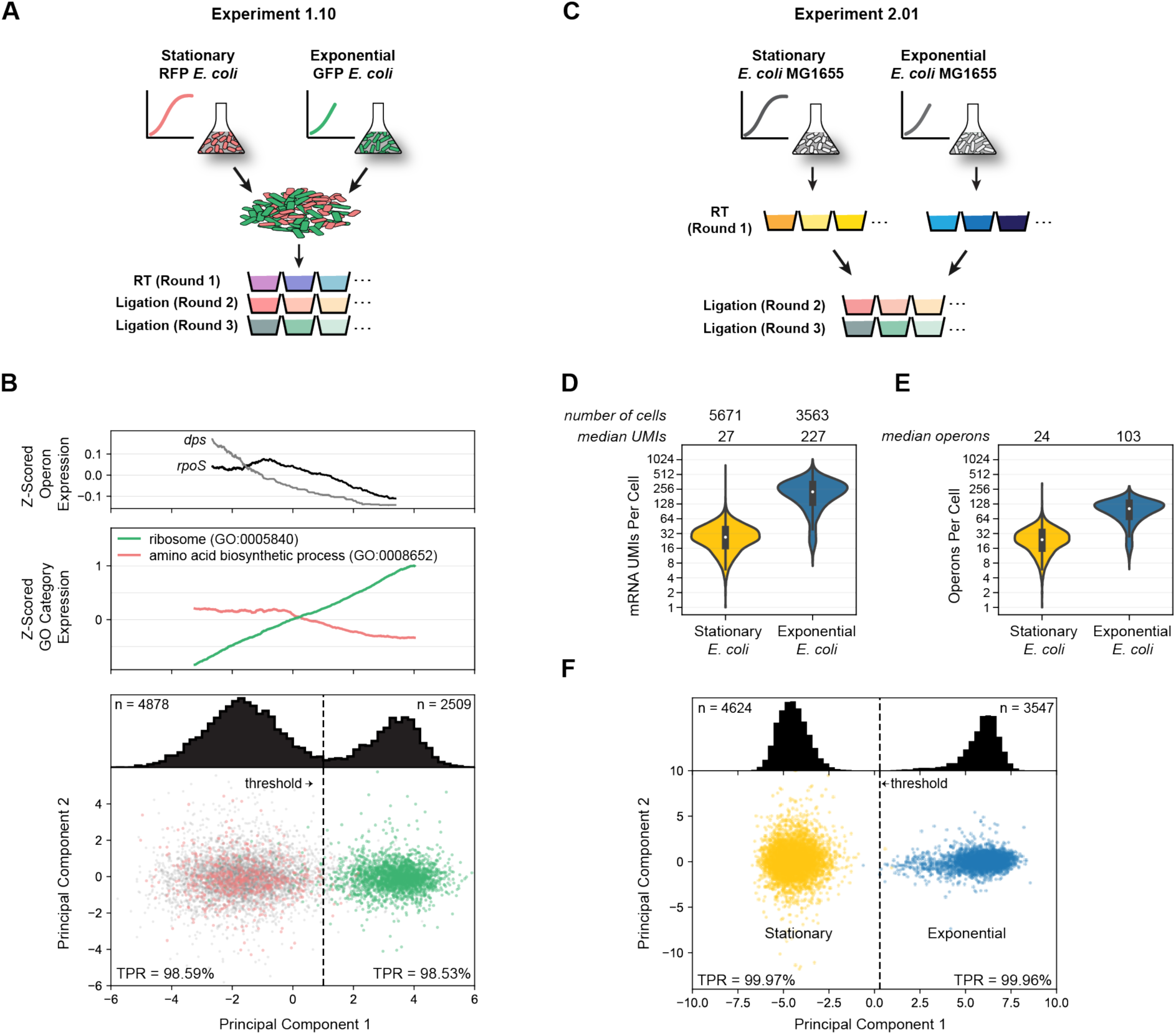
Principal component analysis distinguishes exponential and stationary single *E. coli* cells by mRNA expression patterns (**A**) Schematic of experiment 1.10. Stationary RFP-expressing *E. coli* were mixed with exponential GFP-expressing *E. coli* for barcoding and sequencing by PETRI-seq. Cells were grown separately then combined during fixation. (**B**) *Bottom*: Cells containing UMIs from either the GFP plasmid or the RFP plasmid plotted on PC1 and PC2. The two populations are distinguishable along PC1. Grey points indicate ambiguous cells (no plasmid UMIs). True positive rate (TPR, see calculation in methods) was calculated for RFP cells to the left of the threshold line (PC1 = 1.0) and GFP cells to the right of the threshold line. The TPR refers to the probability that a given cell to the left or right of the threshold line is RFP-expressing stationary or GFP-expressing exponential, respectively. Distribution of all cells across PC1, including cells without any plasmid UMIs, is shown above. 7,387 cells are included (4,878 below threshold and 2,509 above). *Middle:* Expression of GO terms associated with exponential to stationary transition. The moving average (size=1200 cells) of the z-scored expression of operons within the GO term is shown. Expression was z-score transformed for each gene and then for each GO term. Both GO terms are significantly correlated with PC1 prior to calculating moving averages (Spearman rank, p<10^-70^). *Top*: Expression of genes involved in exponential to stationary transition along PC1. The moving average (size=2400 cell) of the z-scored operon expression is shown. Both operons are significantly correlated with PC1 prior to calculating moving averages (Spearman rank, *dps*: p=10^-29^, *rpoS*: p=0.003, FDR<0.01). (**C**) Schematic of Experiment 2.01. Exponential wild-type *E. coli* and stationary wild-type *E. coli* were prepared independently and barcoded separately during round 1 (RT). Exponential *S. aureus* cells were prepared independently and combined with both cell types prior to RT (not shown) for downstream quantification of multiplet frequency and intercellular contamination. (**D**) Distributions of mRNA UMIs captured per stationary or exponential cell after optimized PETRI-seq (2.01). (**E**) Distributions of operons per stationary or exponential cell after optimized PETRI-seq (2.01). Cell numbers are the same as (D). (**F**) Exponential and stationary cells (Experiment 2.01) plotted on PC1 and PC2, which were calculated from these single-cell transcriptomes. Distribution of all cells across PC1 is shown above. Threshold line at PC1 = 0.28 results in a true positive rate (TPR) of 99.97% (stationary, left) or 99.96% (exponential, right). 8,171 cells are included (4,624 below threshold and 3,547 above).

However, we used the PC1 threshold to predict the states of the ambiguous cells and found that 92.2% were stationary cells. Over-representation of stationary cells in the ambiguous population was not surprising as plasmid expression in stationary cells was generally lower than in exponential cells. Importantly, separation of the two transcriptional states was similarly robust in another biological replicate (Fig. S8B) or when operon counts were normalized using sctransform^26^, an alternative method (Fig. S8C). Finally, we investigated expression patterns for operons and gene ontology (GO) terms and found many expected trends related to the transition from exponential growth to stationary phase (Fig. 3B, S8D). For example, *rpoS*, the stationary phase sigma-factor^27^, and *dps*, a DNA-binding protein essential for cellular transition into stationary phase^28^, were upregulated along PC1, as expected in the direction of stationary cells (Fig. 3B, top). Consistent with induction of the stringent response^29^, stationary cells showed a large-scale reduction in ribosomal protein expression as well as an increase in expression of amino acid biosynthetic operons (Fig. 3B, middle).

To further evaluate the power of PETRI-seq to distinguish cells in different growth states, we performed Experiment 2.01 in which exponential and stationary *E. coli* cells were barcoded separately during RT and then pooled for subsequent steps (Fig. 3C). First, by further permeabilizing cells with detergent before ligations and using a higher concentration of ligation primers (Fig. S9), we substantially improved the capture relative to previous experiments (Fig. S6A,D; S8A,B). Specifically, Experiment 2.01 captured a median of 227 and 27 mRNA UMIs per exponential and stationary *E. coli* cell, respectively (Fig. 3D), which corresponded to a median of 103 or 24 operons represented per cell (Fig. 3E). Based on estimates that single exponentially-growing *E. coli* cells contain 2,000-8,000 mRNAs^30–32^, we estimate our capture rate to be roughly 2.5-10% for these cells. For *S. aureus*, we captured a median of 43 mRNA UMIs per cell (Fig. S9E). *S. aureus* cells may contain fewer mRNAs than *E. coli* cells, possibly due to their smaller cell size and genome^33^, though technical differences may also affect capture. We confirmed that optimized PETRI-seq continued to capture single cells with high purity (Fig. S10A-C) comparable to eukaryotic scRNA-seq techniques^23, 24^. Optimized PETRI-seq also captured a high proportion of mRNA relative to rRNA for exponential *E. coli* cells (20.3%, Fig. S9F), though this fraction was reduced for stationary cells (5.5%, Fig. S9G). Lastly, in Experiment 2.01, we again found that PETRI-seq robustly distinguished single *E. coli* cells by growth state (Fig. 3F). Comparison of these sub-populations in Experiments 1.10 and 2.01 corroborated the single-cell purity of PETRI-seq (Fig. S11).

Given ∼20-200 mRNA UMIs per average bacterial cell, we anticipate that future PETRI-seq studies will benefit from aggregation of similar cells in order to define consensus states for sub-populations within heterogenous communities. As a demonstration, we generated consensus transcriptomes by aggregating the expression counts from varying numbers of single cells at either exponential (Fig. S12A,C) or stationary phase (Fig. S12B,D). As expected, correlations with independently prepared bulk libraries from cells in the same growth state increased as more cells were included. Notably, the correlations were stronger and increasing at a greater rate for single-cell/bulk libraries of cells in the same state (colored vs gray curves in Fig. S12C,D), indicating that the aggregated single cells were asymptotically approaching a transcriptome reflecting their growth state.

A key advantage of scRNA-seq over bulk methods is the capacity to characterize rare populations exhibiting distinct gene expression programs. We applied PCA to 6,663 *S. aureus* single-cell transcriptomes generated by PETRI-seq (Fig. S13A) and found that the eight operons most highly correlated with PC1 (Fig. S13B) were lytic genes of prophage ϕSA3usa (Fig. S13C, red arrows)^34, 35^. Cells expressing these operons diverged from the rest of the population along PC1 (Fig. S13A, red points), indicating that PC1 might be capturing rare prophage induction in the *S. aureus* culture. Within the small population, 3 cells exhibited dramatic upregulation of phage lytic transcripts reaching roughly 80% of these single-cell transcriptomes (Fig. S13D). The remaining 25 cells contained fewer than 10% phage transcripts. In further analysis of the heterogeneity in gene expression across the entire *S. aureus* population, we found that for most operons, transcriptional noise^30^ (σ^2^/μ^2^) inversely scaled with mean expression (μ) and followed a Poisson expectation (μ = σ^2^) (Fig. S13E), as was described in other single cell studies^36, 37^. *SAUSA300_1933-1925*, a phage lytic operon encoding putative phage tail and structural genes, clearly diverged and exhibited higher noise than expected from the mean (Fig. S13E), which recapitulated its hypervariability in expression as found by PCA. Similar analysis in *E. coli* discovered a few candidate operons displaying high transcriptional noise (Fig. S14A) that warrant independent validation by methods such as smFISH^36, 38^. One of these, *fimAICDFGH,* encoding type I fimbriae, is known to exhibit population-level phase-variable expression due to promoter inversion^39^. As such, PETRI-seq can detect rare cells occupying distinct transcriptional states and genes displaying high transcriptional heterogeneity within a population.

With a straightforward experimental pipeline requiring no advanced equipment and a per-cell cost of 5.6 cents, PETRI-seq is an efficient and highly affordable method for single-cell RNA sequencing of bacterial populations (Table S1). We sequenced ∼30,000 *E. coli* and *S. aureus* cells with high single-cell purity and found that aggregated transcriptomes from single cells were highly correlated with bulk RNA-seq libraries. PETRI-seq assigned >98% of single cells within isogenic *E. coli* populations to their correct growth phases (i.e. stationary or exponential). Moreover, the high throughput capacity of PETRI-seq was vital for detecting a rare sub-population undergoing prophage induction in 0.04% of *S. aureus* cells. This has important clinical implications, as prophage induction is intimately linked to bacterial pathogenesis^40, 41^.

Optimization of mRNA capture and library preparation is likely to further improve the sensitivity of PETRI-Seq and decrease its cost. Most notably, a customized tagmentation protocol during library preparation could theoretically increase the mRNA capture by 2-fold (see “Future directions for optimization” in Methods). Following our initial deposit of an earlier version of this manuscript on *biorXiv*^42^, and during its formal review, Kuchina and colleagues deposited a manuscript on *biorXiv*^43^ reporting a conceptually similar split-pool based bacterial scRNA-seq method in which *in situ* polyadenylation was utilized to capture mRNAs. It will be of great interest to compare and contrast these methods and to further improve the performance of PETRI-seq. We anticipate that PETRI-seq will be a highly useful tool with broad applications such as characterization of rare, clinically important populations (e.g., persisters^44, 45^) and high-resolution capture of native microbial communities, including unculturable components, a major current challenge in microbiology^46^.

## Methods

### Experimental Methods

#### Bacterial Strains and Growth Conditions

*E. coli* MG1655 was routinely grown in MOPS EZ Rich defined medium (M2105, Teknova, Hollister, CA). pBbE2A-RFP was a gift from Jay Keasling^47^ (Addgene plasmid # 35322). RFP was induced with 20 nM anhydrotetracycline hydrochloride (233131000, Acros Organics, Geel, Belgium). GFP was expressed from p_rplN_-GFP^48^. Plasmid-containing cells were grown in appropriate antibiotics (50 μg/mL kanamycin, 100 μg/mL carbenicillin). *S. aureus* USA300^34^ was routinely grown in trypticase soy broth (TSB) medium (211825, BD, Franklin Lakes, NJ). All bacterial strains were grown at 37°C and shaken at 300 rpm.

#### Custom Primers Used in this Study

All single-tube primers are shown in Table S2. All primer sequences for 96-well split-pool barcoding are shown in Table S3. Primers were purchased from Integrated DNA Technologies (IDT, Coralville, IA).

#### Preparation of Ligation Primers

Concentrations of ligation primers increased for Experiment 2.01, our most optimized version of PETRI-seq. For the information below, quantities are provided as used for Experiment 2.01 (“4x ligation primers”), while alternative quantities are provided in *italics* for libraries using “1x” ligation primers.

Round 2 ligation primers (Table S3) were diluted to 100 μM (*20 μM)*. Round 3 ligation primers were diluted to 70 μM (*20 μM*). Linker SB83 was diluted to 100 μM (*20 μM*). Linker SB80 was diluted to 70 μM (*20 μM*). To anneal barcodes to linkers, 96-well PCR plates (AB0600, Thermo Scientific) were prepared with 3.52 (*4.4*) μL of diluted SB83, 2.64 (*0.8*) μL water, and 3.84 (*4.8*) μL of each Round 2 barcode or 6.6 (*4.4*) μL of diluted SB80, 7.2 (*4.8*) μL of each Round 3 barcode, and 0 (*0.8*) μL water (water only added for “1x” protocol). Primers were annealed by heating the plate to 95°C for 3 minutes then decreasing the temperature to 20°C at a ramp speed of −0.1°C/second.

Primers SB84 and SB81 were also annealed (to form an intramolecular hairpin) prior to blocking by heating 50 μL or 80 μL, respectively, of each 400 (*100*) μM primer to 94°C and slowly reducing the temperature to 25°C.

#### Cell Preparation for PETRI-Seq

For sequencing and qPCR measurements, cells were grown overnight then diluted into fresh media (1:100 for *S. aureus,* p_rplN_-GFP *E. coli*, and wild-type *E. coli*, 1:50 for pBbE2A-RFP *E. coli*) with inducer and antibiotics when applicable. For exponential cells, *E. coli* and *S. aureus* cultures were grown for approximately 2 hours until reaching an OD600 of 0.4 or 0.9, respectively. Exponential *E. coli* cells were used for all qPCR optimization experiments. For stationary cells, pBbE2A-RFP *E. coli* cells were grown an additional 3 hours until the culture reached an OD600 of ∼4 (Experiment 1.06: OD = 4, Experiment 1.10: OD = 3.68). For wild-type stationary cells, *E. coli* cells were diluted 1:100 and grown for ∼3.75 hours to OD600 ∼4 (Experiment 2.01: OD = 3.87). For the combined exponential *E. coli* library (Experiment 1.06SaEc), 3.5 mL of exponential GFP *E. coli* was combined with 3.5 mL of exponential RFP *E. coli*. The *S. aureus* library was prepared separately from 7 mL of exponential cells. For the libraries of exponential GFP *E. coli* combined with stationary RFP *E. coli* (1.06, 1.10), 3 mL of exponential GFP cells was added to ∼300 μL of stationary RFP cells. For Experiment 2.01, 7 mL of exponential wild-type *E. coli* and 7 mL of stationary wild-type were independently fixed. Before fixation, cells were pelleted at 5,525xg for 2 minutes at 4°C. Spent media was removed, and cells were resuspended in 7 mL of ice-cold 4% formaldehyde (F8775, Millipore Sigma, St. Louis, MO) in PBS (P0195, Teknova). This suspension was rotated at 4°C for 16 hours on a Labquake Shaker (415110, Thermo Scientific).

Fixed cells were centrifuged at 5,525xg for 10 minutes at 4°C. The supernatant was removed, and the pellet was resuspended in 7 mL PBS supplemented with 0.01 U/μL SUPERase In RNase Inhibitor (AM2696, Invitrogen, Carlsbad, CA), hereafter referred to as PBS-RI. Cells were centrifuged again at 5525xg for 10 minutes at 4°C then resuspended in 700 μL PBS-RI. Subsequent centrifugations for cell preparation were all carried out at 7000xg for 8-10 minutes at 4°C. Cells were centrifuged, then resuspended in 700 μL 50% ethanol (2716, Decon Labs, King of Prussia, PA) in PBS-RI. Cells were next washed twice with 700 μL PBS-RI, then resuspended in 105 μL of 100 μg/mL lysozyme (90082, Thermo Scientific, Waltham, MA) or 40 μg/mL lysostaphin (LSPN-50, AMBI, Lawrence, NY) in TEL-RI (100 mM Tris pH 8.0 [AM9856, Invitrogen], 50 mM EDTA [AM9261, Invitrogen], 0.1 U/μL SUPERase In RNase inhibitor [10x more than in PBS-RI]). Cells were permeabilized for 15 minutes at room temperature (∼23°C). During optimization of PETRI-seq, we found that the yield from *in situ* RT was not significantly changed by titrating the lysozyme concentration from 4 to 500 μg/mL (data not shown). After permeabilization, cells were centrifuged then washed with 175 μL PBS-RI then resuspended in 175 μL PBS-RI. 100 μL was taken for subsequent steps and centrifuged, while the remaining 75 μL was discarded. Cells were resuspended in 40 μL DNase-RI buffer (4.4 μL 10x reaction buffer, 0.2 μL SUPERase In RNase inhibitor, 35.4 μL water). 4 μL of DNase I (AMPD1, Millipore Sigma) was added, and cells were incubated at room temperature for 30 minutes. To inactivate the DNase I, 4 μL of Stop Solution was added, and cells were heated to 50°C for 10 minutes with shaking at 500 rpm (Multi-Therm, Benchmark Scientific, Sayreville, NJ). 50°C instead of 70°C was used to avoid cell lysis. After DNase inactivation, cells were pelleted, washed twice with 100 μL PBS-RI, then resuspended in 100 μL 0.5x PBS-RI. Cells were counted using a hemocytometer (DHC-S02 or DHC-N01, INCYTO, Chungnam-do, Korea).

#### Split-Pool Barcoding for PETRI-Seq

For RT, Round 1 primers (Table S3) were diluted to 10 μM then 2 μL of each primer was aliquoted across a 96-well PCR plate. A mix was prepared for RT with 240 μL 5x RT buffer, 24 μL dNTPs (N0447L, NEB, Ipswich, MA), 12 μL SUPERase In RNase Inhibitor, and 24 μL Maxima H Minus Reverse Transcriptase (EP0753, Thermo Scientific). 3 * 10^7^ cells were added to this mix. For species-mixed libraries, *E. coli* and *S. aureus* cells were combined at this point. Water was added to bring the volume of the reaction mix to 960 μL. 8 μL of the reaction mix was added to each well of the 96-well plate already containing RT primers, making the final volume in each well 10 μL. The plate was sealed and incubated as follows: 50°C for 10 minutes, 8°C for 12 seconds, 15°C for 45 seconds, 20°C for 45 seconds, 30°C for 30 seconds, 42°C for 6 minutes, 50°C for 16 minutes, 4°C hold. After RT, the 96 reactions were pooled into one tube. At this point, detergent was added for Experiment 2.01, our most optimized version of PETRI-seq. Specifically, 5% Tween-20 was diluted 125x to a final concentration of 0.04%. We measured the volume of the pooled cells to determine this exact volume. Cells were then incubated on ice for 3 minutes, then PBS-RI was added to bring the final concentration of Tween-20 to 0.01% (i.e. add 2508 μL to 836 μL sample, splitting the samples into multiple Eppendorf tubes for centrifugation). Cells were then centrifuged at 10,000xg for 20 minutes at 4°C. The supernatant was removed. For experiments without detergent, cells were centrifuged (10,000xg, 20 minutes, 4°C) immediately after pooling. Without detergent, a cell pellet was not visible after this centrifugation, but with detergent a cell pellet was visible.

For the first ligation, cells were then resuspended in 600 μL 1x T4 ligase buffer (M0202L, NEB). The following additional reagents were added to make a master mix: 7.5 μL water, 37.5 μL 10x T4 ligase buffer, 16.7 μL SUPERase In RNase Inhibitor, 5.6 μL BSA (B14, Thermo Scientific), and 27.9 μL T4 ligase, making the final volume 695.2 μL. 5.76 μL of this mix was added to each well of a 96-well plate containing 2.24 μL of annealed Round 2 ligation primers for a final volume 8 μL. Ligations were carried out for 30 minutes at 37°C. After this incubation, 2 μL of blocking mix (37.5 μL 400 μM SB84 (*100 μM for “1x”*), 37.5 μL 400 μM SB85 (*100 μM*), 25 μL 10x T4 ligase buffer, 150 μL water) was added to each well, and reactions were incubated for an additional 30 minutes at 37°C. Cells were then pooled into a single tube.

The following reagents were added to the pooled cells for the third round of barcoding for Experiment 2.01 (most optimized protocol): 46 μL 10x T4 ligase buffer, 12.65 μL ligase, and 115 μL water. 8.51 μL of this mix was added to each well of a 96-well plate containing 3.49 μL Round 3 ligation primers.

Alternatively, for “1x” primers, the following reagents were added to the pooled cells: 15.6 μL water, 48 μL 10x T4 ligase buffer, and 13.2 μL T4 ligase. 8.64 μL of this mix was added to each well of a 96-well plate containing 3.36 μL of annealed round 3 ligation primers.

The plate was incubated for 30 minutes at 37°C. 10 μL of round 3 blocking mix (72 μL 400 μM SB81 (*100 μM* for “1x”), 72 μL 400 μM SB82 (*100 μM*), 120 μL 10x T4 ligase buffer, 336 μL water, 600 μL 0.5 M EDTA) was added to each well. Cells were then pooled into a single tube. When detergent was used (most optimized protocol), Tween-20 was added to a final concentration of 0.01%. With or without detergent, cells were then centrifuged at 7,000xg for 10 minutes at 4°C. When including detergent, cells were resuspended in 500 μL TEL-RI + 0.01% Tween-20. Without detergent, cells were resuspended in 50 μL TEL-RI (because cell retention is very poor in large volumes without detergent). At this stage, additional washing may be advantageous to reduce any contamination from ambient cDNA (Fig. S5, S10), though we have yet to test this. This suspension was centrifuged at 7000xg for an additional 10 minutes at 4°C, the supernatant was removed, and the cells were resuspended in 30 μL TEL-RI. Cells were counted using a hemocytometer. Aliquots of ∼10,000 cells were taken and diluted in 50 μL lysis buffer (50 mM Tris pH 8.0, 25 mM EDTA, 200 mM NaCl [AM9759, Invitrogen]). 5 μL of 20 mg/mL proteinase K (AM2548, Invitrogen) was added to the cells in lysis buffer. Cells were lysed for 1 hour at 55°C with shaking at 750 rpm (Multi-Therm). Lysates were stored at −80°C.

### Library Preparation for PETRI-Seq

Library preparation steps prior to PCR amplification should be performed with care as any loss of material during these steps result in a reduction in total UMI capture per cell.

Lysates were purified with AMPure XP beads (A63881, Beckman Coulter, Brea, CA) at a 1.8x ratio (∼99 μL). cDNA was eluted in 20 μL water. 14 μL water, 4 μL NEBNext Second Strand Synthesis Reaction Buffer, and 2 μL NEBNext Second Strand Synthesis Enzyme Mix (E6111S, NEB) were added to the purified cDNA. This reaction was incubated at 16°C for 2.5 hours. The resulting double-stranded cDNA was purified with AMPure XP beads at a 1.8x ratio (∼72 μL). cDNA was eluted in 20 μL water and used immediately for tagmentation or stored at −20°C.

cDNA was tagmented and amplified using the Nextera XT DNA Library Preparation Kit (FC-131-1096, Illumina, San Diego, CA). The manufacturer’s protocol was followed with the following modified reagent volumes and primers: 25 μL TD, 20 μL cDNA, 5 μL ATM, 12.5 μL NT, 2.5 μL N70x (Nextera Index Kit v2 Set A, TG-131-2001, Illumina), 2.5 μL i50x (E7600S, NEB), 20 μL water, 37.5 μL NPM. Libraries were amplified for 8 cycles according to the manufacturer’s protocol. After 8 cycles, 5 μL was removed, added to a qPCR mix (0.275 μL EvaGreen [31000, Biotium, Fremont, CA], 0.11 μL ROX Low Reference Dye [KK4602, Kapa Biosystems, Wilmington, MA], 0.115 μL water), and further cycled on a qPCR machine. qPCR amplification was used to determine the exponential phase of amplification, which occurred after 11 cycles for Experiments 1.06SaEc and 1.10 and after 8 cycles for Experiment 2.01. The remaining PCR (not removed for qPCR) was thermocycled an additional 11 or 8 cycles, resulting in a total of 19 or 16 PCR cycles. Products were purified with AMPure XP beads at a 1x ratio and eluted in 30 μL water. The concentration of the library was measured using the Qubit dsDNA HS Assay Kit (Q32854, Invitrogen) and the Agilent Bioanalyzer High Sensitivity DNA kit (5067-4626, Agilent, Santa Clara, CA). Though we used a 1x ratio of AMPure beads for the libraries presented here, we note that, after sequencing, a significant fraction of molecules were too short to be assigned to a BC and/or aligned to the genome (Table S4). A lower ratio of AMPure beads or an additional round of purification might be helpful to reduce the abundance of these wasted reads.

Libraries were sequenced for 75 cycles with the NextSeq 500/550 High Output Kit v2.5 (20024906, Illumina). Cycles were allocated as follows: 58 cycles read 1 (UMI and barcodes), 17 cycles read 2 (cDNA), 8 cycles index 1, 8 cycles index 2.

### Modifications Tested to Optimize PETRI-Seq

To test fixing cells immediately from cultures without centrifugation, ice-cold 5% formaldehyde in PBS was added directly to cells in spent media to bring the final concentration of formaldehyde to 4%. Cell preparation with no lysozyme or no DNase was carried out by simply omitting the enzyme and using water to replace that volume.

Template switching was carried out by adding 2.5 μL 100 μM SB14, 20 μL Maxima H Minus 5x Buffer, 10 μL dNTPs, 2.5 μL SUPERase In RNase Inhibitor, 2 μL Maxima H Minus Reverse Transcriptase, 3 μL water, and 20 μL betaine (J77507VCR, Thermo Scientific) to 40 μL of AMPure purified lysate. SB14 was heated to 72°C for 5 minutes prior to combining the above reagents. The reaction was incubated at 42°C for 90 minutes then heat inactivated at 85°C for 5 minutes. The reaction was purified with AMPure XP beads at a 1.8x ratio and eluted in 30 μL. The purified cDNA was then amplified by setting up the following PCR: 10 μL 5x PrimeSTAR GXL Buffer, 0.1 μL 10 μM SB86, 0.1 μL 10 μM SB15, 1 μL PrimeSTAR GXL Polymerase (R050B, Takara Bio, Kusatsu, Japan), 1 μL dNTPs, and 8 μL water. The reaction was heated to 98°C for 1 minute and then thermocycled 10 times (98°C 10 seconds, 60°C 15 seconds, 68°C 6 minutes). The products were purified by AMPure XP beads at a 1.8x ratio and eluted in 30 μL. The DNA concentration was measured using the Qubit dsDNA HS Assay Kit, and tagmentation was performed according to the manufacturer’s protocol using the appropriate primers (described above for standard PETRI-seq).

For library “1.06SaEc-replicate” (Table S4), we included an “RT clean-up” step as part of library preparation. RT clean-up was carried out in the same way as template switching, but SB14 (TSO) was not added. After incubating the reaction at 42°C for 90 minutes then heat inactivating at 85°C for 5 minutes, reaction components were added for second strand synthesis (70 μL water, 20 μL NEB second strand buffer, 10 μL NEB second strand enzyme). Second strand synthesis was carried out as described and the double-stranded cDNA was used as input for tagmentation. While “RT clean-up” resulted in a broader size distribution on the bioanalyzer after tagmentation (not shown), it did not change the yield of PETRI-seq and thus was not used for other libraries.

For library 1.08 (Table S4), we included a longer RT (∼2 hours) using the following thermocycling protocol: 50°C 10 min, 10x: 8°C 12s, 15°C 45s, 20°C 45s, 25°C 5 min, 42°C 1 min, 50°C 2 min.

### qPCR Quantification After *In Situ* DNase or *In Situ RT*

For qPCR quantification after *in situ* RT, cells were counted prior to RT, and then the *in situ* RT reaction described above (scaled to one 50 μL reaction) was set up with equal cell numbers for each condition and technical replicate. A random hexamer (SB94) or a gene-specific primer (SB10) was used as an RT primer. After RT, cells were centrifuged at 7,000xg for 10 minutes then washed in 50 μL PBS-RI. After one wash, cells were resuspended in 50 μL lysis buffer, and 5 μL of proteinase K was added. Cells were lysed for 1 hour at 55°C with shaking at 750rpm. For qPCR quantification after *in situ* DNase treatment, cells were washed twice after DNase treatment, as described for PETRI-seq cell preparation, then lysed.

Unpurified lysates were diluted 50x (except for ethanol vs. no ethanol, which were diluted 10x) in water and heated to 95°C for 10 minutes to inactivate proteinase K. Diluted lysates were then used directly in qPCR with either Kapa 2x MasterMix Universal (KK4602, Kapa Biosystems) or *Power* SYBR Green Master Mix (4368706, Applied Biosystems, Foster City, CA). For quantification of genomic DNA after DNase treatment or quantification of cDNA after RT with random hexamers, qPCR primers SB5 and SB6 were used, and relative abundances were calculated based on an experimentally determined amplification efficiency of 88%, which corresponded to an amplification factor of 1.88. Relative abundance thus referred to 1.88^-ΔCt^, where ΔC_t_ was the difference between the C_t_ value of each sample and a calibrator C_t_. For RT with the gene-specific primer, qPCR primers SB12 and SB13 were used, as SB12 anneals to the gene-specific primer (SB10). The experimentally determined amplification factor for these primers was 1.73. To quantify cDNA yield, the abundance of a matched sample with no RT (processed equivalently but RT enzyme omitted) was subtracted from each measurement. All replicates were technical replicates, which were treated independently during and after the condition tested.

### qPCR Quantification of Ligation Efficiency

To test barcode ligation with a 16-base linker relative to a 30-base linker, approximately 1 μg of purified RNA (bulk) was used for RT with either SB110 or SB114 (used as a positive control). RT was carried out as described for *in situ* RT, scaled to 50 μL. cDNA was then purified with AMPure XP beads. SB113, the primer to be ligated, was annealed either to SB111 (30 bases) or SB83 (16 bases). 2.24 μL of the annealed primers was then used in a 10 μL ligation reaction. The products were purified with AMPure XP beads. To quantify the proportion of ligated product, qPCR was performed with SB86 and SB13, which amplifies only the ligated product, as SB86 anneals to the ligated overhang, or SB115 and SB13, which amplifies all RT product, as SB115 anneals to the RT primer overhang. ΔΔC_t_ was calculated for the two primer sets with RT product from SB114 as a reference [ΔΔC_t_ = ΔC_t_(experimental, ligated) - ΔC_t_(control, SB114 RT), ΔC_t_ = C_t_(SB86,SB13) - C_t_(SB115,SB13)]. SB114 includes primer sites for both SB86 and SB115, so it mimics ligation with 100% efficiency.

### Test of DNase Inactivation by Incubating Cells with Exogenous DNA

After DNase treatment, inactivation, and two PBS-RI washes (described above), cells were resuspended in 20 μL PBS-RI. 6 μL was removed and added to 1 μL DNase reaction buffer, 1 μL water, and 2 μL of a 775 bp PCR product (800 ng). As a control, 1 μL DNase I was added instead of 1 μL water. The reactions were incubated for 1 hour, after which 1 μL of stop solution was added. The cells were centrifuged for 10 minutes at 7,000xg. The supernatants were then heated to 70C for 10 minutes to inactivate DNase. 5 μL of each reaction was run on a gel.

### Bulk Library Preparation

For preparation of bulk samples from fixed cells (shown in Fig. 3D, 3E, S12), 25 μL (∼10^7^ cells) was taken after PETRI-seq cell preparation and just prior to *in situ* RT. These cells were centrifuged and resuspended in 50 μL lysis buffer supplemented with 5 μL proteinase K. Cells were lysed at 55°C for 1 hour with shaking at 750 rpm (Multi-Therm). RNA was then purified from lysates with the Norgen Total RNA Purification Plus Kit (48300, Norgen Biotek, Ontario, Canada). 300 μL buffer RL was added to the lysate before proceeding to the total RNA purification protocol. Alternatively, the standard bulk RNA sample (shown in Figure S3B) was prepared by centrifuging a cell culture at 5525xg for 2 minutes at 4°C then resuspending cells in 1mL of PBS-RNAprotect (333 μL RNAprotect Bacteria Reagent [76506, Qiagen, Hilden, Germany], 666 μL PBS). For immediate RNA stabilization by RNAprotect (Fig. S3B), 2 mL of RNAprotect was immediately added to 1 mL of exponential *E. coli* cells. For immediate RNA stabilization by flash freezing, 330 μL 60% glycerol was added to 1 mL exponential *E. coli* cells, and cells were flash frozen in ethanol and dry ice (<1 minute). Frozen cells were put at −80°C overnight, then thawed, spun down and re-suspended in PBS-RNAprotect. For all three protocols, after resuspending cells in RNAprotect, cells were then pelleted again, and RNA was prepared with the Norgen Total RNA Purification Plus Kit according to the manufacturer’s instructions for Gram-negative bacteria.

Purified RNA from either protocol was treated with DNase I in a 50 μL reaction consisting of 2-5 μg RNA, 5 μL DNase Reaction Buffer, 5 μL DNase, and water. Reactions were incubated at room temperature for 30-40 minutes. Reactions were purified by adding 300 μL buffer RL and proceeding according to the Norgen total RNA purification protocol. Total RNA was depleted of rRNA using the Gram-Negative Ribo-Zero rRNA Removal Kit (MRZGN126, Illumina), purified by ethanol precipitation, and resuspended in 10 μL water. For RT, 6 μL RNA was combined with 4 μL Maxima H Minus 5x Buffer, 2 μL dNTPs, 0.5 μL SUPERase In RNase Inhibitor, 1 μL SB94, 0.5 μL Maxima H Minus Reverse Transcriptase, 4 μL betaine, and 2 μL water. The reaction was thermocycled as follows: 50°C for 10 minutes, 8°C for 12 seconds, 15°C for 45 seconds, 20°C for 45 seconds, 30°C for 30 seconds, 42°C for 6 minutes, 50°C for 16 minutes, 85°C 5 minutes, 4°C hold. For second strand synthesis, 14 μL water, 4 μL NEBNext Second Strand Synthesis Reaction Buffer, and 2 μL NEBNext Second Strand Synthesis Enzyme Mix were added directly to the RT mix. This reaction was incubated at 16°C for 2.5 hours. Double-stranded cDNA was purified with AMPure XP beads at a 1.8x ratio (∼72 μL beads) and eluted in 30 μL water. Purified cDNA was used for tagmentation with the Nextera XT kit according to the manufacturer’s protocol. Bulk libraries were purified twice with AMPure XP beads at a 0.9x ratio. The resulting libraries were quantified and sequenced as described for PETRI-seq libraries above.

### Growth Curves

Overnight cultures were grown as described above and then diluted 1:100 into 1 mL EZ Rich Defined Media with or without 20 nM aTc. Antibiotics were added for plasmid-containing strains. For each condition, 100 μL of diluted cells were aliquoted into 4 wells of a 96-well plate. The plate was incubated at 37°C with shaking on the plate reader (Synergy Mx, Biotek, Winooski, VT). OD600, GFP, and RFP were measured every 10 minutes.

## Computational Methods

### Barcode Demultiplexing, Cell Selection and Alignment

*Cutadapt*^49^ was used to trim low-quality read 1 and read 2 sequences with phred score below ten. *Umi_tools*^50^ was used in paired-end mode to extract the seven base UMI sequence from the beginning of read 1. Read pairs were then grouped based on their three barcode sequences using the *cutadapt* demultiplex feature. FASTQ files were first demultiplexed by barcode 3, requiring that matching sequences were anchored at the beginning of the read, overlapped at 21 positions (“--overlap 21”, including downstream linker [GGTCCTTGGCTTCGC]), and had no more than 1 mismatch relative to the barcode assignment (-e 0.05). As part of demultiplexing, the barcode and linker sequence were trimmed in read 1. For barcode 2, *cutadapt* was used to locate barcode sequences with the expected downstream linker, allowing no more than 1 mismatch (-e 0.05 --overlap 20) and requiring the barcode at the beginning of the read. The barcode and linker sequences were trimmed. Next, reads were demultiplexed by barcode 1, requiring the barcode at the beginning of the read and allowing 1 mismatch but no indels. The final output after demultiplexing was a set of read 1 and read 2 FASTQ files where each file corresponded to a three-barcode combination (BC). The “knee” method^5^ was used to identify BCs for further processing. Briefly, each BC was sorted by descending total number of reads, and then the cumulative fraction of reads for each BC was plotted. Because the yield per BC could be better assessed later after collapsing reads to UMIs, an inclusive threshold was used at this stage to select BCs for downstream processing, which allowed for more precise cell selection after downstream processing (Fig. S4C). *Cutadapt* was then used to trim and discard read 2 sequences containing barcode 1 or the linker sequence. Note that at this point all necessary information was contained in the read 2 FASTQ files, so further processing did not consider the read 1 files. Next, cDNA sequences were aligned to reference genomes using the backtrack algorithm in the Burrows-Wheeler Alignment tool, *bwa*^51^, allowing a maximum edit distance of 1 for assigned alignments.

### Annotating Features and Grouping PCR Duplicates by Shared UMI

*FeatureCounts*^52^ was used to annotate operons based on the alignment position. Operon sequences were obtained from RegulonDB^53^ and ProOpDB^54^ for *E. coli* and *S. aureus*, respectively. Because *featureCounts* uses an “XT” sam file tag for annotation, the *bwa* “XT” tag was first removed from all sam files using a python script. The resulting bam files after *featureCounts* were used as input for the group function of *umi_tools* with the “--per-gene” option in directional mode^50^. The directional algorithm is a network-based method that identifies clusters of connected UMI sequences to group as single UMIs. The result was a set of bam files with UMI sequences corrected based on probable errors from sequencing or amplification. A python script was used to collapse reads to UMIs. Reads with the same BC, error corrected UMI, and operon assignment were grouped into a single count. With 4^7^ possible UMIs, we confirmed that the expected rate of UMI collisions (different molecules with the same UMI) was low by implementing a correction based on the Poisson expectation of collisions^55^. As this correction had a negligible effect, we did not include it for other analysis. Reads mapping to multiple optimal positions were omitted except rRNA alignments for which multiple alignments were expected. The distribution of number of reads per UMI for all UMI-BC-operon combinations was plotted to establish a threshold below which UMIs were excluded (Fig. S4D). Filtered UMIs were used to generate an operon by BC count matrix. Anti-sense transcripts were removed. BCs with fewer than a threshold of total UMIs were then removed (Fig. S4F,G and Fig. S9H,I).

### Bulk Sequencing Libraries

For bulk sequencing libraries, only read 2 was used for alignment in order to mimic single-cell methods. Bulk sequencing libraries were pre-processed to remove adapters using *cutadapt*^49^. *Trimmomatic*^56^ was then used to remove leading or trailing bases below quality phred33 quality 3 and discard reads shorter than 14 bases. Surviving reads were aligned using the backtrack algorithm in *bwa* ^51^ with a maximum edit distance of 1. Reads with more than one optimal alignment position were removed. *FeatureCounts*^52^ was used to generate a matrix of operon counts for the bulk libraries. To compare single-cell libraries generated by PETRI-seq to bulk samples, the UMI counts for a given set of BCs (e.g. GFP-expressing *E. coli*) were summed for all operons. A count matrix was then generated as described for bulk libraries. To calculate TPM, raw counts were divided by the length of the operon in kilobases. Then, each length-adjusted count was divided by the sum of all adjusted counts divided by 1 million.

### Calculating Multiplet Frequency

The multiplet frequency was defined as the fraction of non-empty BCs corresponding to more than one cell. To calculate the predicted multiplet frequency, the proportion of predicted BCs with 0 cells was calculated based on a Poisson process: 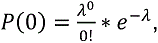 the proportion of BCs with 1 cell was calculated: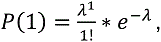 and the proportion with greater than 1 cell was calculated: *P(≥ 2) = 1 − P(1) − P(0)*. Finally, the multiplet frequency was calculated: 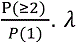 was the fraction of cells relative to total possible BCs – for example,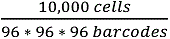 = 0.011 = λ. The experimental multiplet frequency was computed from the species-mixing experiment as described for populations with unequal representation of two species^20^.

### Principal Component Analysis (PCA)

rRNA and all plasmid genes (RFP, GFP, AmpR, KanR, tetR) were first removed from the count matrix. Operons with 5 or fewer total counts in the library were also removed (except for Figure S7 in which all operons with >0 counts were included). Cells with fewer than 15 mRNAs were removed. Total operon counts for each cell were normalized by dividing each count by the total number of counts for that cell then multiplying the resulting value by the geometric mean^13^ of the total mRNA counts for each cell. The scaled values were then log transformed after adding a pseudocount to each. For each operon, expression values were scaled to z-scores^57^. Principal components were computed using scikit-learn in python.

To normalize counts using sctransform in Seurat^26^, first rRNA and all plasmid genes were removed from the count matrix. Operons with 10 or fewer total counts, and cells with fewer than 15 mRNAs were also removed. A Seurat object was created in R from the resulting matrix, and sctransform was applied. The resulting scaled counts were used as input for PCA.

True positive rate (TPR) was calculated as follows, using red cells to the left of a threshold line as an example: 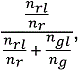 where *n^rl^* = number of red cells left of threshold, *n^r^* = total number of red cells, *n^gl^* = number of green cells left of threshold, *n^g^* = total number of green cells.

### Computing Moving Averages of Gene Expression Along PC1

Using a custom Python script, the cells in the normalized, log-transformed, z-scored gene matrix were sorted by PC1. The rolling function in the pandas package was then used to compute rolling averages of the size indicated for each figure. Win_type was set to “None”. The corresponding PC1 coordinate was the moving average of the PC1 values. Moving averages for GO terms were computed as described, except the z-scored sum of z-scored counts for all operons in the GO term was used to calculate the moving average instead of expression from a single operon. In cases where multiple genes from the same operon were included in a GO term, only one gene was included. Significance of expression trends was determined by the Spearman rank correlation between the operon or GO term expression and PC1, prior to calculating a moving average. FDR was determined by the Benjamini-Hochberg procedure^58^.

### Computing Operon Noise

Noise was defined as σ^2^/μ^2^, where σ is standard deviation and μ is mean. Noise and mean were calculated for all operons with at least 5 raw counts (UMIs) in the dataset (either *S. aureus* or *E. coli*). Count matrices were normalized by cell and multiplied by the geometric mean of total UMIs per cell in the library (but not log-transformed) before computing noise and mean. Operons with mean expression < 0.002 after normalization were excluded. To calculate a p-value for the divergence of SAUSA300_1933-1925, a line was fit to the log-scaled noise vs log-scaled mean of the data. The residuals of the experimental data to the best-fit line were calculated and z-scored. The p-value was determined based on a normal distribution of the z-scored residuals. For the *E. coli* dataset, cells with BC2 #22, 49, or 69 were removed because in rare cases these barcodes misaligned to an operon, resulting in the appearance of hypervariability in gene expression.

### Future directions for optimization

We anticipate the following modifications would further improve the final mRNA capture of PETRI-seq. During the library preparation step of PETRI-seq, subjecting double-stranded cDNA to conventional tagmentation with both N5 and N7 adaptors (Illumina Nextera XT) reduces the mRNA capture by ∼ 2-fold. This is because only one of the adaptors (N7 in our case) could be subsequently amplified, leading to the loss of all molecules tagmented by N5. Thus, modified tagmentation using, for example, a commercially available and customizable Tn5 (Lucigen) could potentially increase capture by 2-fold^13^.

Secondly, capture may be improved by further increasing primer and enzyme concentrations during the ligation steps and/or using a hairpin ligation^13^ instead of an inter-molecular linker. For instance, increasing the concentration of round 3 ligation primers by 4-fold alone increased mRNA capture by 2.7-fold in both exponential and stationary *E. coli* cells (Fig. S9A). Our preliminary results also indicate that adding polyethylene glycol (PEG) to the third round of ligations increases capture up to 1.3x (not shown).

Given that rRNAs comprise >95% of total RNA species in many bacteria, we reason that mRNA capture might be additionally improved by designing RT primers with sequences biased against rRNA^59^, thereby directing reagents preferentially toward mRNA. Alternatively, *in situ* 5’-phosphate-dependent exonuclease treatment could be used to preferentially degrade processed RNAs, the majority of these being rRNAs^60^, prior to RT.

As the capture rate of PETRI-seq increases, it will become more important to reduce the fraction of reads mapped to rRNA in order to saturate read coverage of mRNAs in the library. For this purpose, abundance-based normalization by melting and rehybridization of double-stranded cDNA followed by duplex-specific nuclease treatment^61^ may also be considered to deplete dsDNAs encoding rRNAs.

In addition to the capture rate, further reduction in cost and time will improve the PETRI-seq workflow. We have preliminary results indicating that DNase treatment may not be necessary (not shown). However, we have not yet determined if omitting the DNase buffer incubation or heat inactivation would alter cell permeability. Without DNase treatment, cell preparation time would be reduced by ∼1.5 hours.

Finally, we have shown that in Experiment 2.01, ∼1-5% of UMIs within a single-cell transcriptome are likely derived from other cells (Fig. S10C). This cross-contamination, which may be the result of ambient cDNA released from cells during or after barcoding, might be reduced by more thorough cell washing prior to lysis. Cross-contamination may also be reduced by preparing lysates with fewer cells, thereby reducing the likelihood of barcode collisions with ambient cDNA (or other cells). PCR may also be a source of cross-contamination through chimera formation or priming by residual barcodes. This type of contamination may be reduced by thorough washing prior to lysis (to remove free barcodes) or by optimizing the parameters of the PCR. Computationally, we also showed that a more stringent alignment reduces the level of apparent cross-contamination resulting from incorrect alignment (Figure S5E,F), but more stringent alignment results in a decrease in captured UMIs per cell (Figure S5C,D,G,H). Future studies could use longer reads (i.e., 150-cycle Illumina Nextseq) to eliminate ambiguities in alignment without sacrificing capture rate.

## Data Availability

Raw data has been submitted to the Gene Expression Omnibus (GEO). An accession number will be included once assigned. All figures except Figures 1, S1, S2 include original data. An overview of all of the experiments can be found in Table S4.

## Code Availability

Relevant code for this manuscript is available upon request.

## Authors’ contributions

SB, WJ, and ST conceived the study. SB, WJ performed experiments and data analysis. PO assisted with computational analysis. SB, WJ, and ST wrote the paper.

## Acknowledgements

We thank the Tavazoie laboratory for helpful discussions and comments on early drafts of the manuscript. ST is supported by award 5R01AI077562 from NIH. SB is supported by NSF award DGE - 1644869. WJ is supported by a fellowship from the Jane Coffin Childs Fund.

## Supplementary Materials

**Figure S1:**
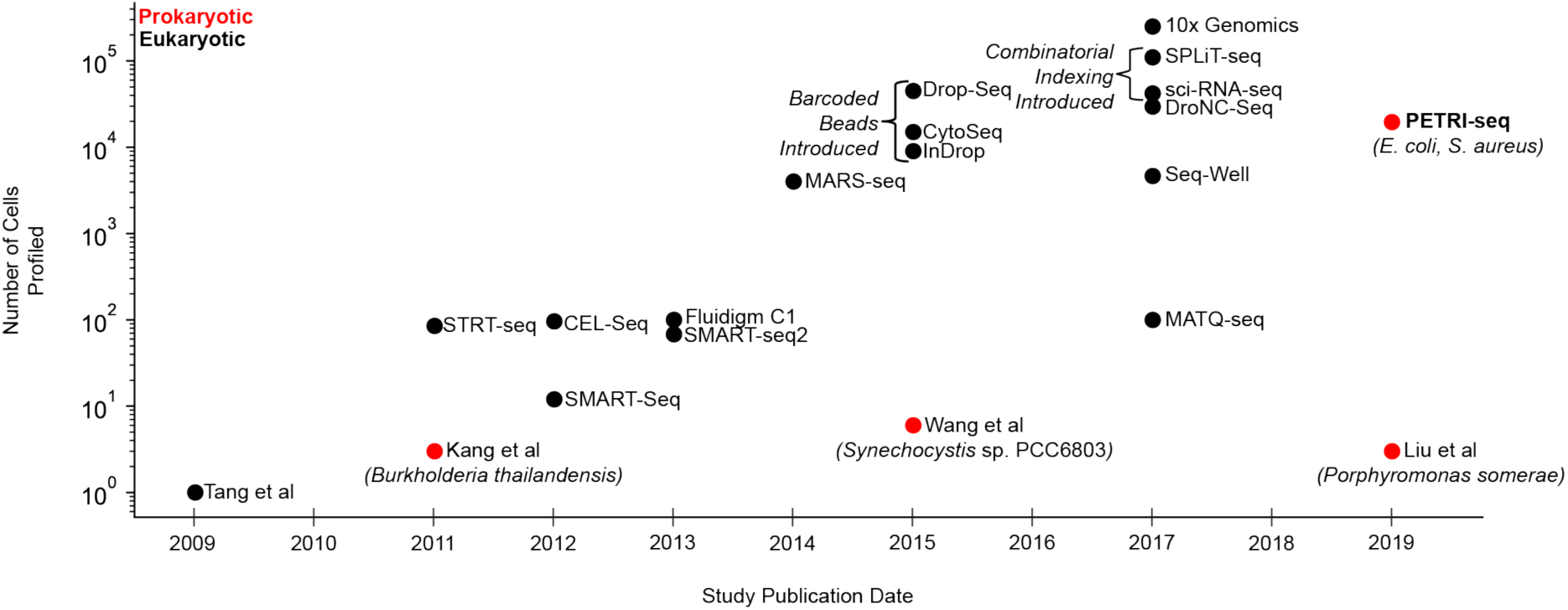
Timeline of scRNA-seq method development Timeline of key developments in eukaryotic^1-6,8,10-12^,^62–69^ and prokaryotic^14–16^ scRNA-seq detailing the number of cells sequenced in each experiment. The eukaryotic timeline is adapted from reference 62. Prokaryotic scRNA-seq has lagged significantly behind eukaryotic scRNA-seq due to technical challenges. In 2011, the first single-cell microarray study by Kang et al was described for a few *Burkholderia thailandensis* cells^14^, each containing 2 pg of RNA, orders of magnitude more than many bacterial species of interest^70^. More recent reports described sequencing of six *Synechocystis* sp. PCC6803 cells^15^ (Wang et al) and three *Porphyromonas somerae* cells^16^ (Liu et al), each of which contains 1-5 fg of RNA. These methods comprehensively characterized the transcriptomes of a few single cells. However, they are prone to contamination and not equipped to study highly heterogeneous bacterial communities and rare populations like persisters^71^ across thousands of cells.

**Figure S2:**
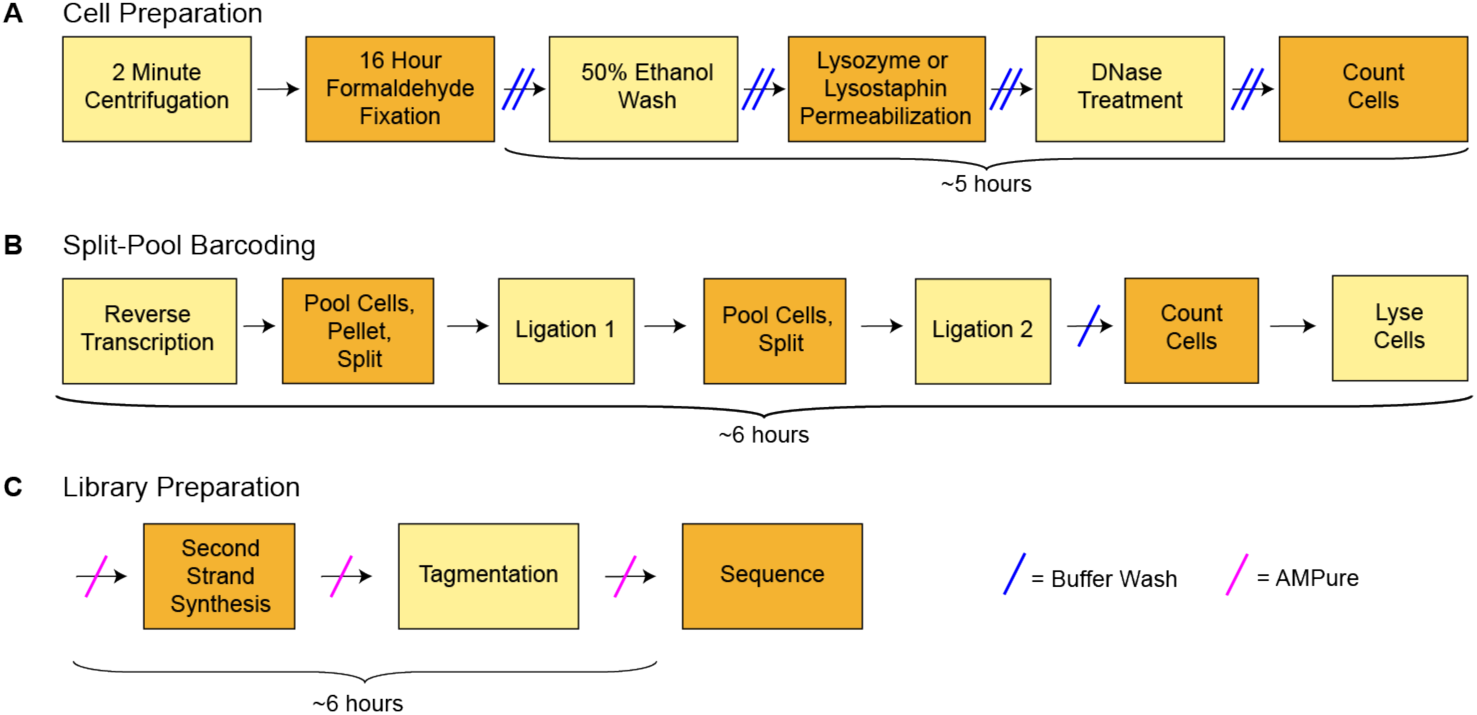
Detailed Schematic of PETRI-seq. PETRI-seq libraries can be prepared in just 2.5 days. (**A**) Detailed schematic of steps for cell preparation, which is started at the end of day 1 and finished on day 2. (**B**) Detailed schematic of steps for split-pool barcoding, which is entirely done on day 2. (**C**) Detailed schematic of steps for library preparation, which can be completed (up to sequencing) on day 3 (or later, if preferred).

**Figure S3:**
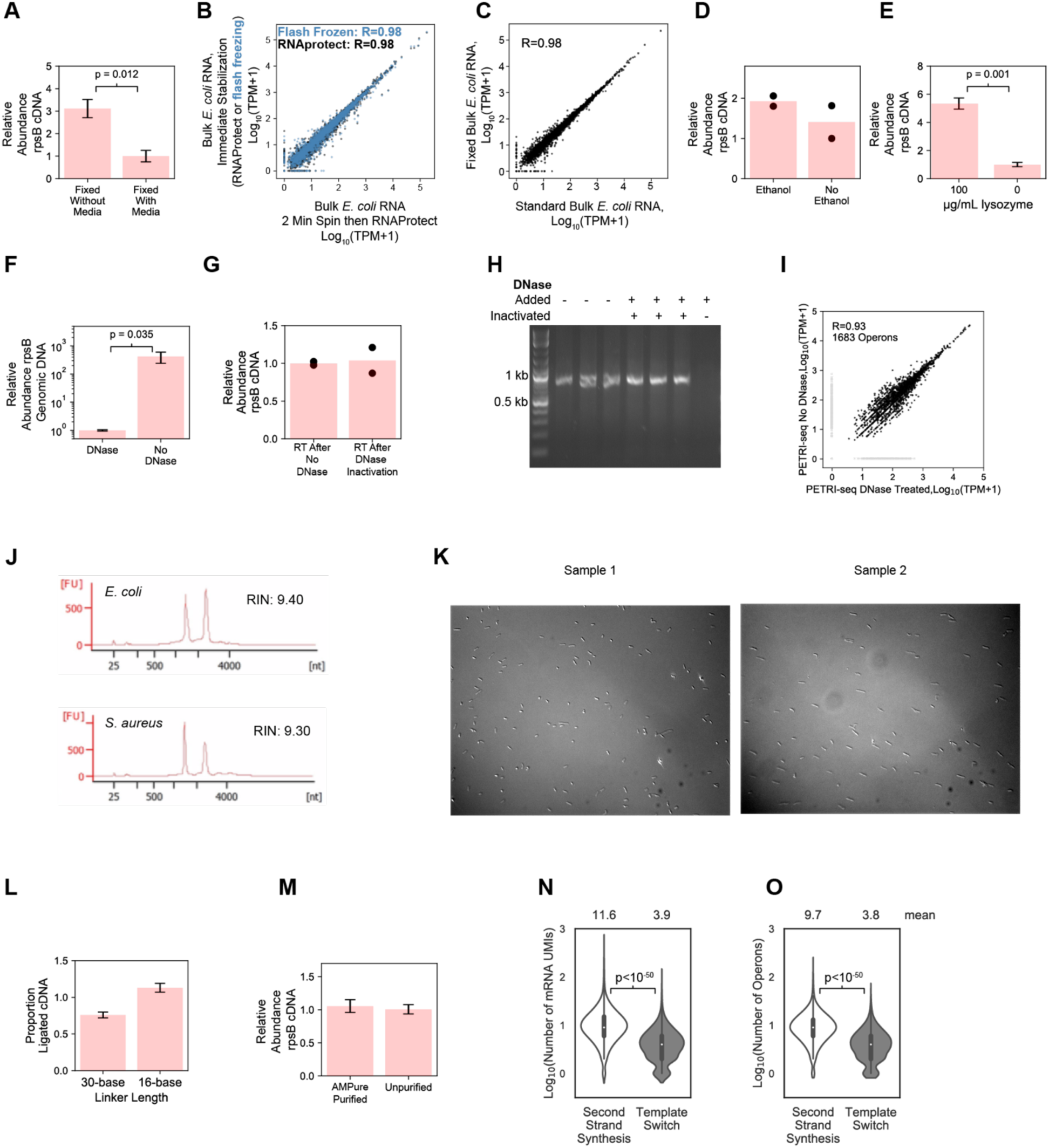
Development and preliminary optimization of PETRI-seq. (**A**) Fixation without media (brief pelleting before fixation) resulted in a higher yield (n=3, p=0.012, 2-sided t-test) of *rpsB* cDNA than fixation with media (formaldehyde added directly to culture). qPCR was done after *in situ* RT with random hexamers. (**B**) 2-minute spin before RNA purification did not alter the bulk transcriptome. Transcriptomes stabilized by RNAprotect after 2-minute spin (x-axis) was compared to ones that were stabilized immediately by either RNAprotect or flash freezing (y-axis). 2,617 operons are included, and Pearson’s r is reported. (**C**) Fixation did not alter the *E. coli* transcriptome. Correlation is shown between RNA purified from *E. coli* cells after being fixed by 4% formaldehyde for 16 hours (“Fixed Bulk”) and RNA purified directly from growing cells (“Standard Bulk”). For both libraries, reverse transcription was done after purifying the RNA. 2,617 operons are included, and Pearon’s r is reported. (**D**) We did not detect a significant change in yield of *rpsB* cDNA when cells were resuspended in 50% ethanol as part of cell preparation (n=2, p=0.35, 2-sided t-test). qPCR was done after cell preparation and *in situ* RT with gene-specific RT primer (SB10) (n=2) (**E**) Lysozyme treatment significantly improved the yield of rpsB cDNA (n=3, p=0.001, 2-sided t-test). qPCR was done after cell preparation and *in situ* RT with random hexamers. (**F**) qPCR after DNase treatment or incubation with DNase buffer only (“No DNase”) confirmed the efficacy of *in situ* DNase treatment (n=8, p=0.035, 2-sided t-test). (**G**) qPCR after cell preparation and RT with gene-specific *rpsB* RT primer (SB10) confirmed that DNase was inactivated, as we did not detect a significant change in the yield of *rpsB* cDNA with or without DNase treatment (n=2, p=0.84, 2-sided t-test). (**H**) Gel of 775-bp PCR fragment after 1-hour incubation with cells prepared for *in situ* RT confirmed inactivation of DNase. For the lane that was not inactivated, DNase was directly added to the incubation of cells with the PCR product. (**I**) DNase treatment did not significantly alter *E. coli* transcriptomes. Aggregated PETRI-seq UMIs from one library treated with DNase and one library not treated with DNase were used to calculate TPM. Both libraries used cells from the same exponential *E. coli* culture. (**J**) After DNase treatment and prior to RT, an aliquot of cells was lysed and column-purified. RNA integrity was assessed by bioanalyzer (DNA HS), which reported high RNA integrity number (RIN) for both *E. coli* and *S. aureus*. (**K**) Microscope images after cell preparation of *E. coli*. (**L**) qPCR after bulk RT and ligation with a 16-base or 30-base linker confirmed that ligation was effective with a 16-base linker. We detected a mild increase in ligation efficiency with the 16-base linker (p=0.001, n=3, 2-sided t-test), though the fold-change between the conditions was minor (1.5x). ΔΔCt was calculated for ligated product relative to total RT product and normalized to cDNA prepared with an RT primer including the ligated sequence (SB114). (**M**) qPCR after cell preparation and *in situ* RT showed that cDNA was retained after AMPure purification (n=4, p=0.69, 2-sided t-test). (**N,O**) Number of mRNA UMIs (K) or operons (J) per BC after PETRI-seq with second strand synthesis or template switch used for library preparation. Second-strand synthesis resulted in significantly more mRNAs per cell (p < 10^-50^, 2-sided Mann-Whitney U) and operons per cell (p < 10^-50^, 2-sided Mann-Whitney U). Libraries contained ∼10,000 BCs and are shown here before further downstream filtering for likely single cells. These were prepared using an unoptimized protocol (Experiment 1.08).

**Figure S4:**
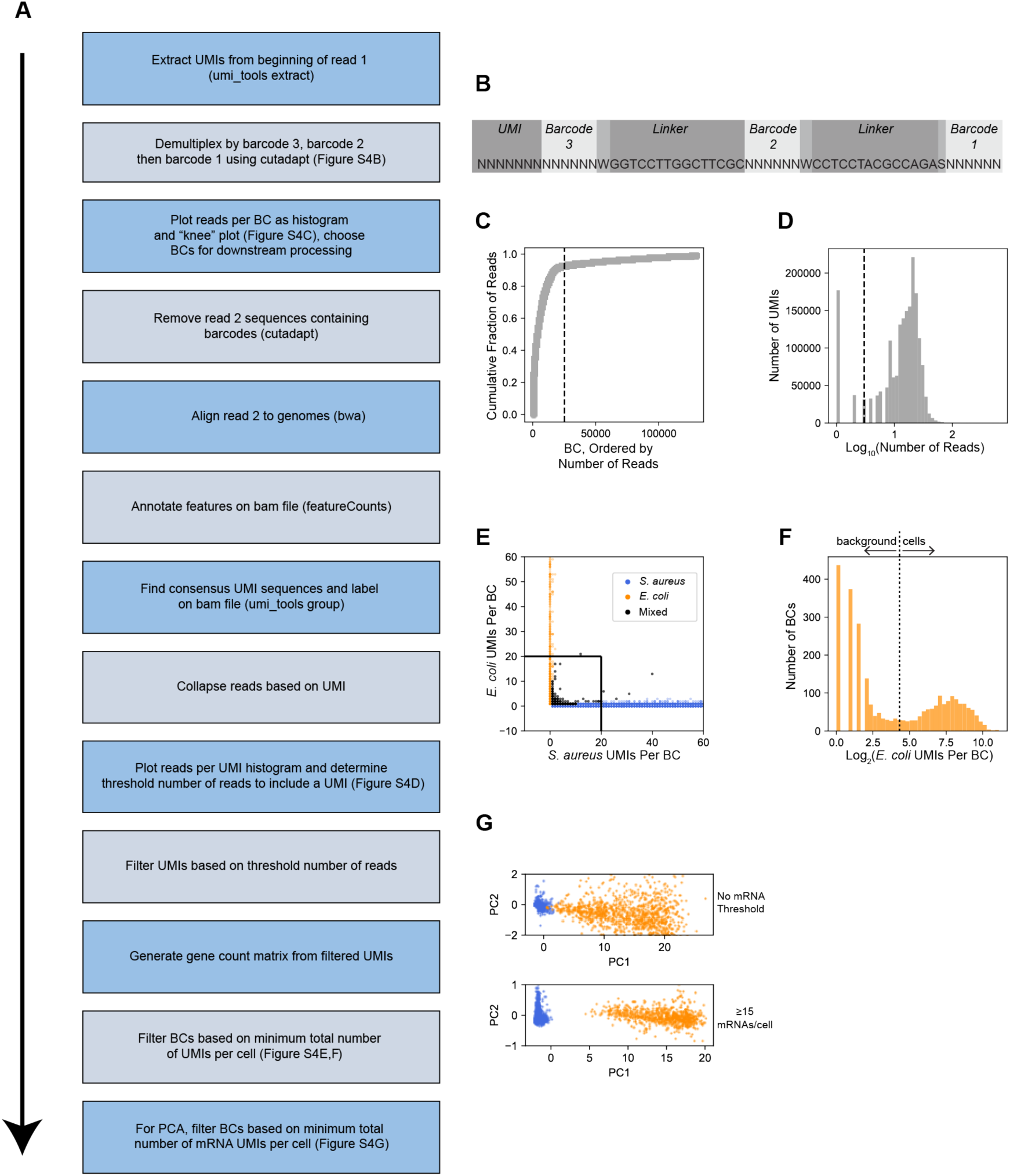
Computational pipeline for PETRI-seq. (**A**) Schematic of computational steps, which are further detailed in Methods. (**B**) Structure of contig elements in read 1 after Illumina sequencing of PETRI-seq. To reduce the length of the sequence, barcodes overlap by one base with the adjacent linker sequence. (**C**) Representative “knee plot” used to select BCs for further analysis. The threshold line at 25,000 BCs is inclusive to facilitate additional filtering after collapsing PCR duplicates to UMIs. (**D**) Representative histogram of reads per UMI. A threshold line was set for each library. For this library, only UMIs with more than 3 reads were kept for downstream analysis. Threshold line at log_10_(3). (**E**) Species mixing plot without all BCs containing >0 UMIs for library 1.06SaEc. BCs with fewer than 20 UMIs per cell were removed from further analysis. Line segments at x=20 and y=20. (**F**) Distribution of *E. coli* BCs from species mixing plot in (D). BCs above the threshold line were used for further analysis and considered single *E. coli* cells. Threshold line at log_2_(20). (**G**) PCAs of *E. coli* (orange) and *S. aureus* (blue) BCs from library 1.06SaEc. For calculation of principal components, rRNA operons were omitted and counts were normalized and scaled as described in methods. *Top*: all *S. aureus* and *E. coli* BCs with greater than 20 total UMIs and greater than 0 mRNAs are included (13,786 *S. aureus*, 1,153 *E. coli*). *Bottom*: Only BCs with greater than or equal to 15 mRNA UMIs are included (6,683 *S. aureus*, 800 *E. coli*). For 100% of *S. aureus* BCs, PC1<0.05, and for 100% of *E. coli* BCs, PC1>4.

**Figure S5:**
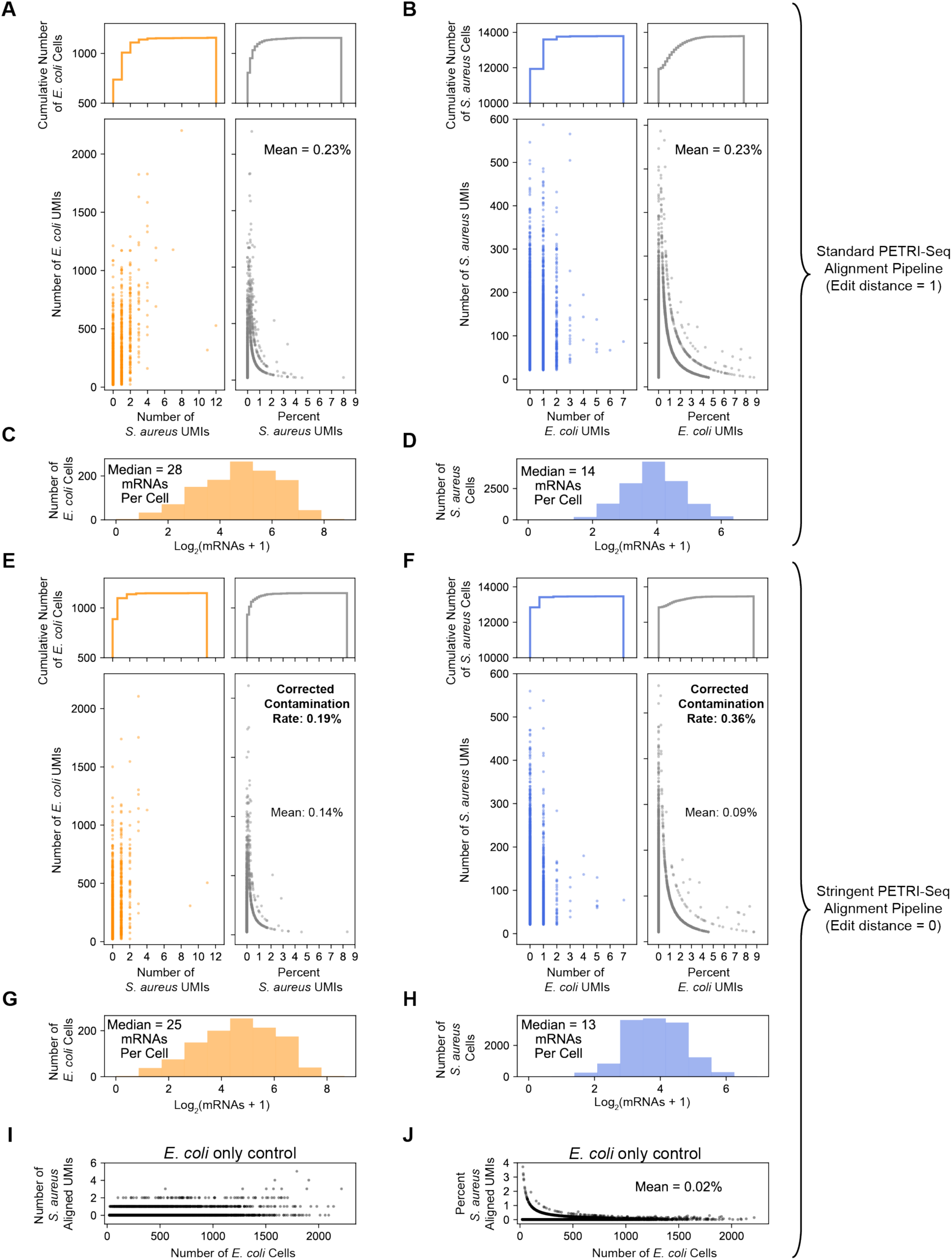
Quantification of intercellular contamination using *E. coli* and *S. aureus* cells After defining single *E. coli* and *S. aureus* cells based on thresholds described in Figure 2B, we further examined levels of cross-contamination within likely single-cells using the mixed species library 1.06SaEc (Table S4), as detailed in the following panels: (**A**) Quantification of *S. aureus-*aligned UMIs assigned to *E. coli* cells after standard PETRI-seq alignment and operon assignment. Standard alignment allows an edit distance of 1 between a 17-bp read and the genome. Reads mapping equally well to *E. coli* and *S. aureus* are discarded. *Bottom left*: Scatterplot of total *E. coli* UMIs and *S. aureus* UMIs assigned to each *E. coli* cell (as defined in Figure 2B). *Bottom right:* Scatterplot of total *E. coli* UMIs per *E. coli* cell and the percent of total UMIs for each cell aligned to *S. aureus.* To summarize these data points, *E. coli* cells include a mean of 0.23% *S. aureus* UMIs. *Top*: Cumulative distributions of *E. coli* cells corresponding to scatterplots below. (**B**) Quantification of *E. coli-*aligned UMIs assigned to *S. aureus* cells after standard PETRI-seq alignment and operon assignment. *Bottom left*: Scatterplot of total *E. coli* UMIs and *S. aureus* UMIs assigned to each *S. aureus* cell (as defined in Figure 2B). *Bottom right:* Scatterplot of total *S. aureus* UMIs per *S. aureus* cell and the percent of total UMIs for each cell aligned to *E. coli. S. aureus* cells include a mean of 0.23% *E. coli* UMIs. *Top*: Cumulative distributions of *E. coli* cells corresponding to scatterplots below. (**C**) Distribution of mRNAs per *E. coli* cell shown in (A). (**D**) Distribution of mRNAs per *S. aureus* cell shown in (B). (**E-H**) To better understand the source of the minor contamination observed in A and B, we repeated the analysis using a more stringent alignment in which an edit distance of 0 was required for a read to be mapped. (**E**) Quantification of *S. aureus* aligned UMIs assigned to *E. coli* cells after stringent PETRI-seq alignment and operon assignment. Reads mapping equally well to *E. coli* and *S. aureus* are discarded. Panels are the same as (A). (**F)** Quantification of *E. coli* aligned UMIs assigned to *S. aureus* cells after stringent PETRI-seq alignment and operon assignment. Panels are the same as (B). (**G**) Distribution of mRNAs per *E. coli* cell shown in (E). As a result of stringent alignment, the number of mRNAs per cell is reduced. (**H**) Distribution of mRNAs per *S. aureus* cell shown in (F). (**I-J**) To further understand the impact of alignment on apparent cross-contamination, we used stringent alignment (edit distance = 0) to map UMIs for a library of only *E. coli* (Experiment 1.10). (**I**) Quantification of UMIs assigned to *S. aureus* after stringent alignment and operon assignment for a PETRI-seq library prepared with only *E. coli*. *S. aureus* UMIs are thus computational artifacts due to misalignment to the *E. coli* genome. (**J**) Quantification of percent of total UMIs assigned to *S. aureus* in library prepared with only *E. coli* after stringent alignment. *E. coli* cells include a mean of 0.02% *S. aureus* aligned UMIs, indicating that the majority of interspecies contamination observed in (E) is due to contamination during PETRI-seq barcoding or library preparation rather than incorrect alignment. To quantify this contamination, we assume conservatively that interspecies contamination in (E) and (F) is entirely due to impurities during PETRI-seq rather than incorrect alignment. Because in this library 25% of UMIs aligned to *E. coli* and 75% of UMIs aligned to *S. aureus*, we needed to correct the percentages of inter-species alignment (0.14% for *E. coli* or 0.09% for *S. aureus*) to predict the percent of UMIs in a given single-cell derived from any other cell (whether or not it is of the same species). We thus predict that 0.19-0.36% of UMIs in a given single-cell transcriptome are derived from a different cell 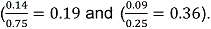 This is reported as the “corrected contamination rate” in E and F.

**Figure S6:**
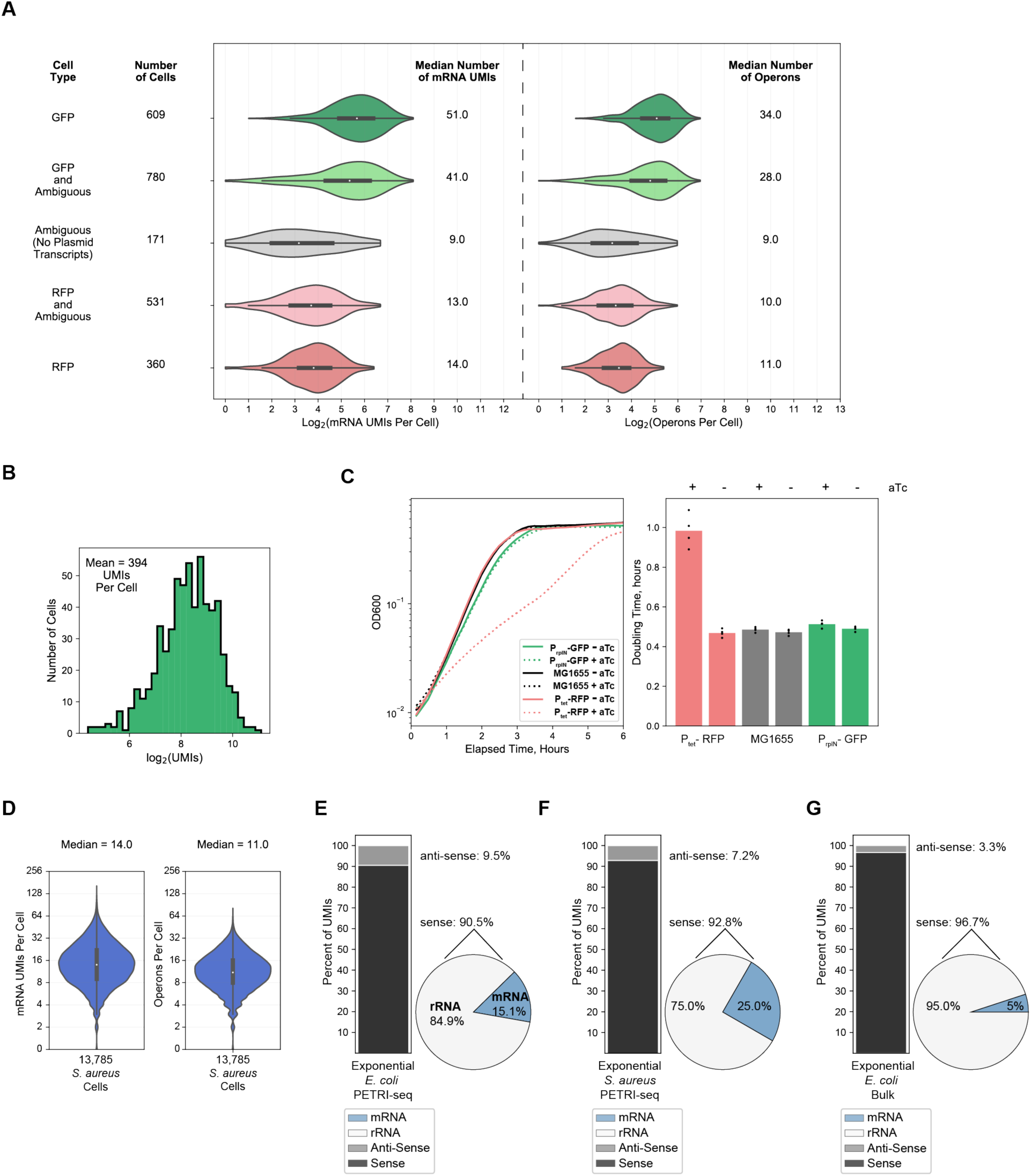
Further evaluation of PETRI-Seq for *E. coli* and *S. aureus* in Experiment 1.06SaEc (A) Distributions of mRNA UMIs (*left*) and operons (*right*) per *E. coli* cell in five sub-populations, including GFP cells (contain GFP plasmid transcripts), RFP cells (contain RFP plasmid transcripts), ambiguous cells (contain no plasmid transcripts), and either RFP or GFP *and* ambiguous cells. Three ambiguous cells classified as *E. coli* in Figure 2B were omitted as they contained zero mRNAs. (B) Distribution of total RNAs per GFP-containing exponential *E. coli* cell. 609 cells are included. (**C**) *Left*, growth curves for PrplN-GFP, Ptet-RFP, and MG1655 (no plasmid) cells with and without aTc. *Right*, doubling times calculated from the growth curves. Ptet-RFP had a significantly longer doubling time than all other strains/conditions when induced with aTc (n=4, p<10-4, 2-sided t-test), which might explain fewer mRNA UMIs in these cells. (**D**) Distributions of mRNA UMIs (*left*) and operons (*right*) per *S. aureus* cell. 13,785 cells are included. 2 cells were omitted as they contained zero mRNAs. (**E,F,G**) Breakdown of total aligned UMIs (E,F) or reads (G) per cell for PETRI-seq exponential GFP- and RFP-expressing *E. coli* (E), PETRI-seq exponential *S. aureus* (F), and bulk exponential wild-type *E. coli* (G). *Left:* Stacked bar shows breakdown of sense and anti-sense alignments. *Right*: Pie shows breakdown of rRNA and mRNA alignments within the sense fraction.

**Figure S7:**
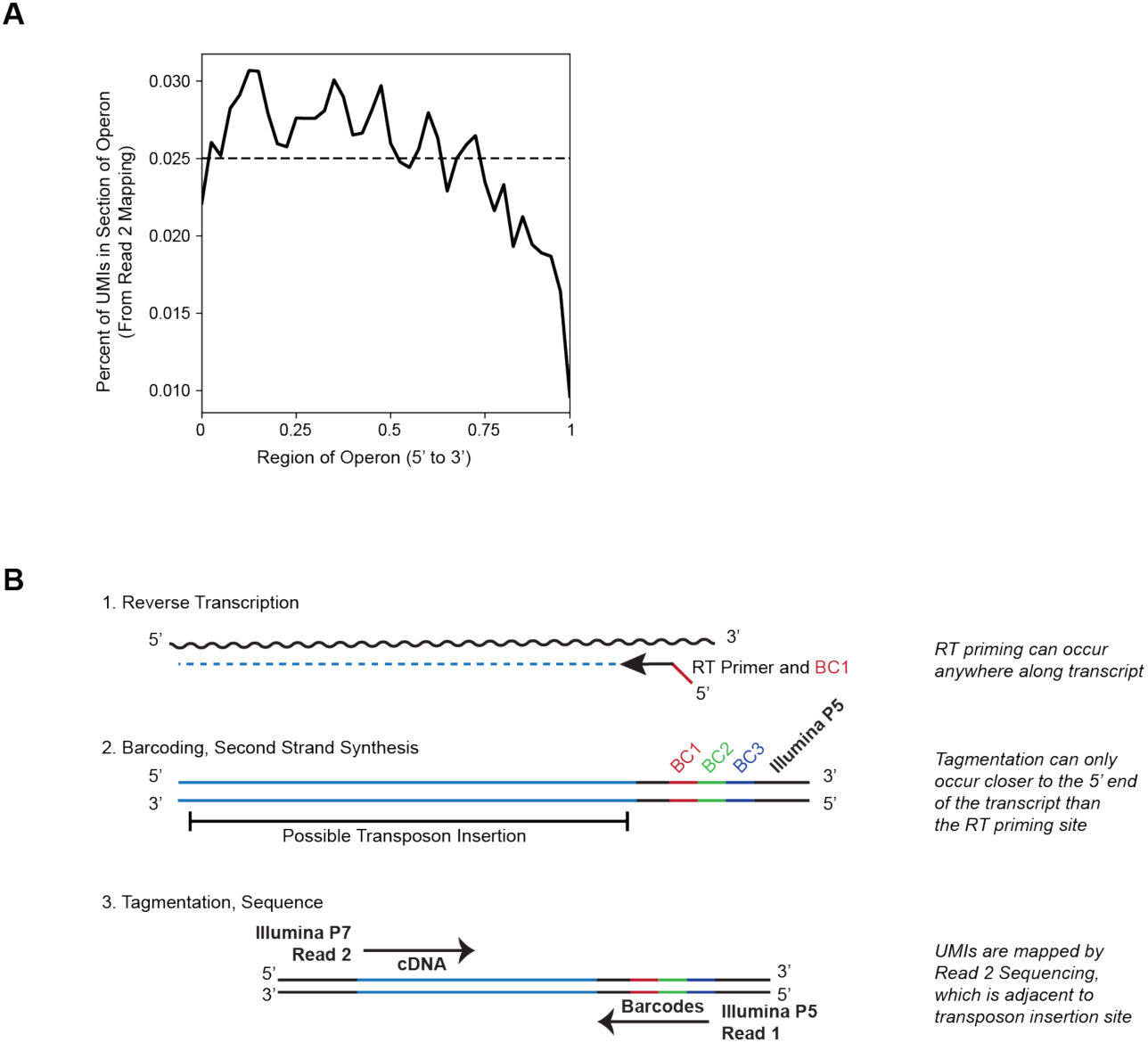
Distribution of UMI alignments along operons in PETRI-seq. (**A**) UMI coverage along *E. coli* mRNA operons. Operons were split into 40 equal bins, and mRNA UMIs aligned within each bin were counted. Horizontal line at 0.025% indicates the expectation if all UMIs were evenly distributed. (**B**) It is clear from the distribution in (A) that PETRI-seq exhibits a mild bias against the 3’ end of mRNA operons. This can partly be explained by the nature of sample preparation in PETRI-seq, which is illustrated here. Random hexamers can prime anywhere along a transcript during RT (*top*), but then tagmentation must occur downstream of the priming site, which is towards the 5’ end of the transcript (*middle*). Reads are then mapped using read 2, which is the sequence adjacent to the site of tagmentation (*bottom*). It therefore is expected that fewer UMIs would map towards the 3’ end of the transcript. We also considered that the bias observed in (A) could result from mRNA degradation during cell preparation, but degradation of RNA before RT appears minimal (Figure S3J).

**Figure S8:**
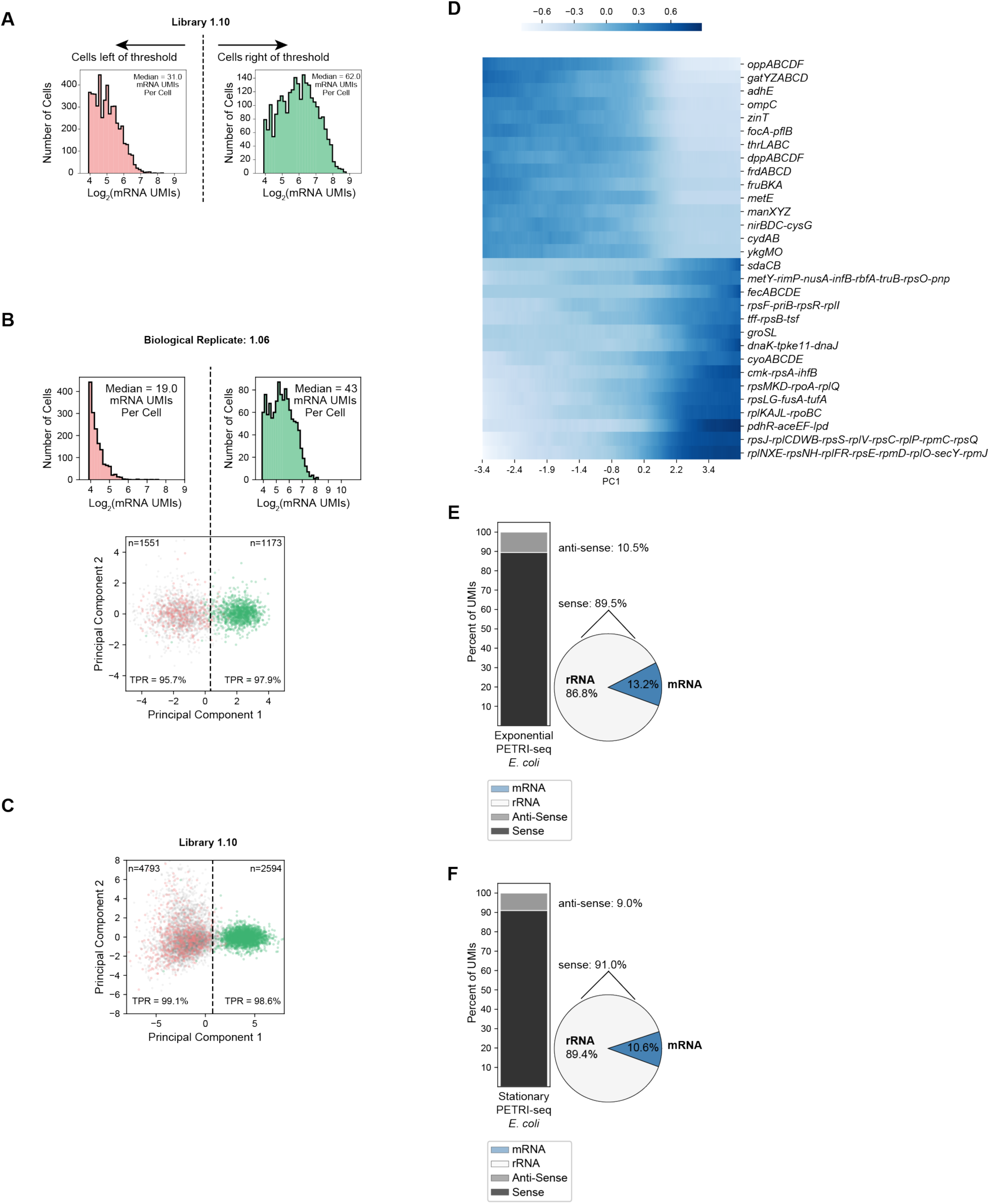
Further evaluation of growth phase characterization by PETRI-Seq. (**A**) Distribution of mRNA UMIs per cell (Experiment 1.10) on either side of the threshold line in Figure 3B. Grey cells (without plasmid UMIs) are included. Only cells with greater than 14 mRNA UMIs per cell were included, as cells with fewer were excluded from the PCA. 4,878 cells are left of the threshold, and 2,509 cells are right of the threshold. (**B**) Biological replicate library (1.06) shows that PETRI-seq can reproducibly distinguish between stationary and exponential cells by projecting cells onto the principal components calculated from the first library (*bottom*). 2,724 cells are included. 1,551 cells are left of the threshold, and 1,173 cells are right of the threshold. mRNA UMIs captured per cell on either side of the threshold line are shown (*top*). (**C**) PCA as in Figure 3B, but UMI counts were normalized using sctransform^26^. (**D**) Expression along PC1 (Figure 3B, Experiment 1.10) of operons with the most positive or negative PC1 loadings (z-scored moving average, size=1,000 cells). (**E,F**) Breakdown of total aligned UMIs per cell for Experiment 1.10 for cells above and below the PC1 threshold in Figure 3B. In (E), Exponential *E. coli* (above the threshold) are shown and in (F), stationary *E. coli* (below the threshold) are shown. *Left:* Stacked bar shows breakdown of sense and anti-sense alignments. *Right*: Pie shows breakdown of rRNA and mRNA alignments within the sense fraction.

**Figure S9:**
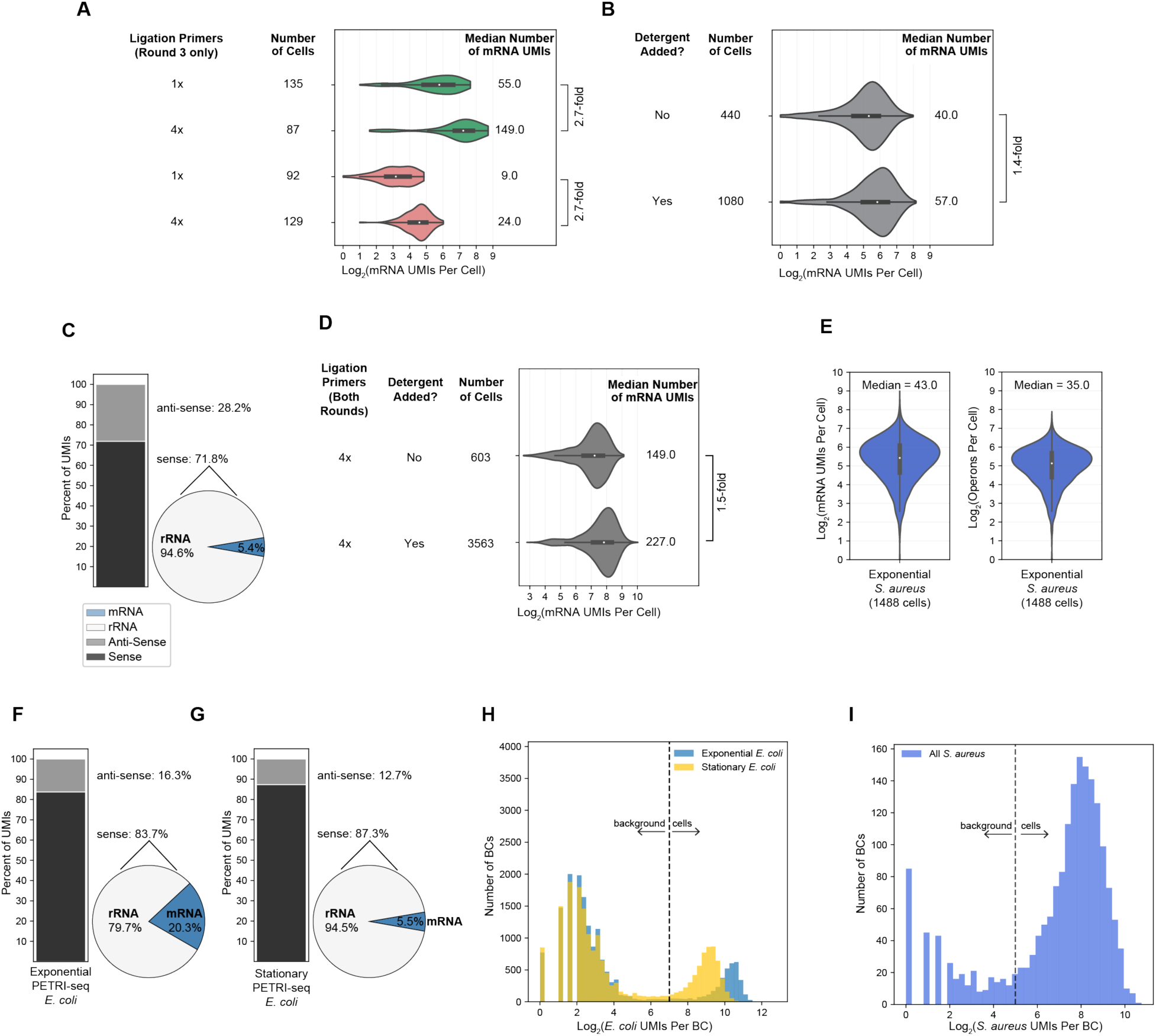
Additional optimization of PETRI-seq by increasing ligation primer concentration and adding detergent during barcoding. (**A**) Increasing the concentration of round 3 ligation primers by 4x relative to original PETRI-seq (Experiments 1.06SaEc and 1.10) increases mRNA UMIs per cell 2.7-fold for GFP-expressing exponential *E. coli* cells (green) and RFP-expressing stationary *E. coli* cells (red). (**B**) Adding detergent (tween-20) to cells after RT and after ligation 3 increased mRNA UMIs per cell 1.4-fold relative to original PETRI-seq for wild-type exponential *E. coli* cells. (**C**) In an alternative PETRI-seq, we used 10x more concentrated RT primers relative to original PETRI-seq. We observed a shift in the breakdown of sense/anti-sense and mRNA/rRNA UMIs with this modification. *Left*: Stacked bar shows breakdown of sense and anti-sense alignments. *Right*: Pie shows breakdown of rRNA and mRNA alignments within the sense fraction. The percent of anti-sense RNAs is significantly increased, while the percent of mRNAs is significantly decreased. Based on these observations, we hypothesized that any condition which effectively increases the intracellular concentration of RT primers could lead to this undesirable shift. For this reason, when detergent was added, it was only ever added after RT to avoid further permeabilizing cells and increasing the effective concentration of RT primer. (**D**) Combining detergent treatment and increased ligation primer (for both ligation rounds, unlike in A) resulted in higher mRNA capture for wild-type exponential *E. coli* cells. Detergent treatment again increased mRNA UMIs per cell (1.5-fold). (**E**) Optimized PETRI-seq with increased ligation primer concentrations and detergent treatment resulted in *S. aureus* transcriptomes with a median of 43 mRNA UMIs per cell (*left*) and 35 operons per cell (*right*). (**F,G**) Breakdown of total aligned UMIs per cell for optimized PETRI-seq (Experiment 2.01) for exponential *E. coli* (F) and stationary *E. coli* (G) *Left:* Stacked bar shows breakdown of sense and anti-sense alignments. *Right*: Pie shows breakdown of rRNA and mRNA alignments within the sense fraction. (**H,I**) Distributions of total UMIs per *E. coli* (H) and *S. aureus* (I) BCs in Experiment 2.01. With more UMIs captured per BC in Experiment 2.01 relative to previous experiments, we imposed higher thresholds for distinguishing cells from background than used previously (Figure S4F). For *E. coli*, BCs with more than 128 total UMIs were considered cells (threshold line in H), while for *S. aureus*, BCs with more than 32 total UMIs (threshold line in I) were considered cells.

**Figure S10:**
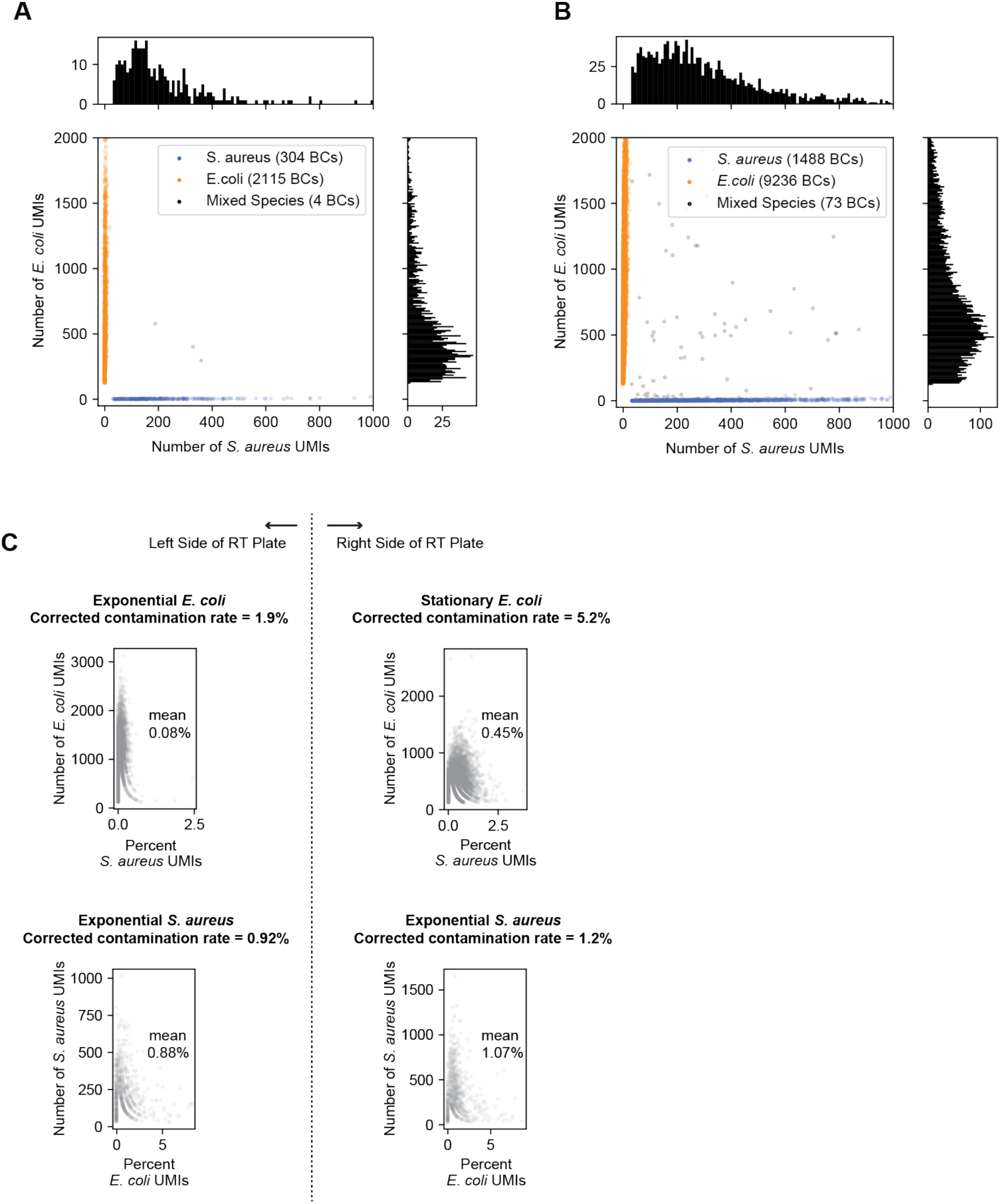
Multiplet frequency and intercellular contamination for optimized PETRI-seq. (**A**) Species mixing plot for PETRI-seq with 4x ligation primers and no detergent. The multiplet frequency is 0.7%, which is 5-fold higher than the Poisson expectation of 0.14% for 2,423 BCs. *E. coli* BCs with > 128 total UMIs and *S. aureus* BCs with > 32 total UMIs were included. (**B**) Species mixing plot for PETRI-seq with 4x ligation primers and detergent (Experiment 2.01). The multiplet frequency is 2.8%, which is 4.7-fold higher than the Poisson expectation of 0.6% for 10,797 BCs. This indicates that compared to no detergent, detergent treatment did not significantly increase multiplet frequency relative to the Poisson expectation. *E. coli* BCs with > 128 total UMIs and *S. aureus* BCs with > 32 total UMIs were included. (**C**) Scatterplots of the percent of total UMIs for each cell aligned to the incorrect species for optimized PETRI-seq (Experiment 2.01). Reads were aligned using the stringent alignment described in Figure S5. *Top left:* Percent of *S. aureus* UMIs in exponential *E. coli* cells (based on first round barcode). Cells included a mean of 0.08% *S. aureus* UMIs. *Top right:* Percent of *S. aureus* UMIs in stationary *E. coli* cells (based on first round barcode). Cells included a mean of 0.45% *S. aureus* UMIs. *Bottom left:* Percent of *E. coli* UMIs in *S. aureus* cells barcoded with exponential *E. coli* (based on first round barcode). Cells included a mean of 0.88% *E. coli* UMIs. *Bottom right:* Percent of *E. coli* UMIs per *S. aureus* cell barcoded with stationary *E. coli* (based on first round barcode). Cells included a mean of 1.07% *E. coli* UMIs. As described in Figure S5, we used these inter- species contamination rates to predict a corrected contamination rate (including intra-species contamination). For experiment 2.01, this results in the following: 1.9% for exponential *E. coli*, 5.2% for stationary *E. coli*, 0.92% for *S. aureus* (left side), and 1.2% for *S. aureus* (right side). Though higher than the contamination rates observed in the previous species mixing experiment (Figure S5E,F), these rates are comparable to common eukaryotic scRNA-seq methods^23, 24^. Furthermore, we anticipate that contamination could be reduced by additional washing prior to cell lysis (see “Future directions for optimization” in Methods). We did not observe a significant change in the corrected contamination library when detergent was not used (cells in panel A, data not shown).

**Figure S11:**
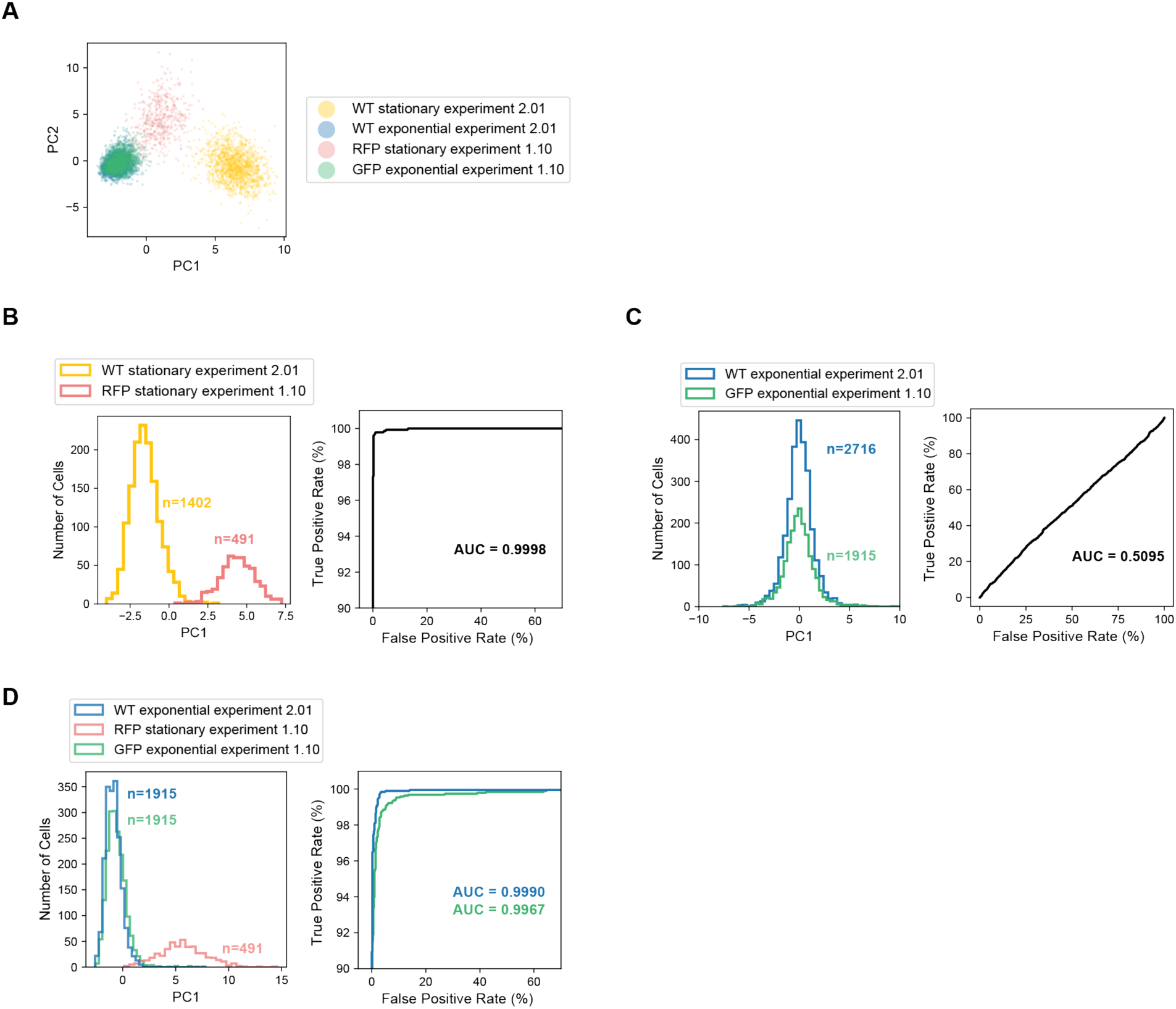
Comparison of plasmid-labeled (Experiment 1.10) and RT-labeled (Experiment 2.01) mixed growth stage libraries reveals minimal cross-contamination between *E. coli* cells barcoded together Comparison of the single-cell transcriptomes from Experiments 1.10 (barcoded together) and 2.01 (barcoded separately) allows for an assessment of cross-contamination between cells in different growth states. Impurities in PETRI-seq can result from multiplets (cells traveling together during barcoding) or more subtle cross-contamination (addressed in Figure S5), which we speculate may come from barcoding of free molecules (not contained within cells) or occasional non-specific addition of barcodes once cells are pooled. In Experiment 2.01, exponential and stationary cells were prepared separately and then barcoded independently during RT. They were then pooled for additional barcoding by ligations, but all UMIs can almost unambiguously be distinguished as originating from stationary or exponential cells based on the barcode added during RT. While we note there is some possibility of RT occurring after the cells are pooled or chimeras forming during PCR and introducing cross-contamination between the populations, we consider the populations of stationary and exponential cells from Experiment 2.01 to be representative of highly pure sampling of each growth stage. In contrast, the RFP-expressing stationary cells and GFP-expressing exponential cells barcoded in Experiment 1.10 were combined for fixation and barcoded entirely together, resulting in more opportunity for cross-contamination. In order to quantify this cross-contamination, we sought to use Experiment 2.01 as a reference. To account for differences in the capture efficiency for the two experiments, we first down-sampled every cell to 30 mRNA UMIs. In (**A**), we show a PCA for all 4 cell types, which reveals that the two stationary populations are biologically distinct. The stationary cells may be distinct because they were grown on different days to slightly different ODs, and RFP cells were induced and grown with aTc, which has a strong effect on growth rate (Fig. S6C). In contrast, the two exponential populations appear very similar (overlapping blue and green populations). We further quantified these findings in B and C. In (**B**), we recalculated PC1 using only the stationary cells from both experiments. *Right*: The receiver operating characteristic (ROC) shows that PC1 is a strong classifier of the two states, thereby establishing that these stationary populations are biologically distinct. In (**C**), we calculated PC1 using only the exponential cells from both experiments. *Right*: The ROC shows that PC1 is a weak classifier of the two exponential states with performance similar to random assignment (Area Under the ROC Curve [AUC]=0.5). (**D**) PC1 was recalculated using wild-type exponential cells from Experiment 2.01, GFP-expressing exponential cells from Experiment 1.10, and RFP-expressing stationary cells from Experiment 1.10 in order to quantify cross-contamination between the GFP and RFP cells using the wild-type exponential cells from experiment 2.01 as a reference. *Right*: ROC shows that PC1 is a strong classifier of exponential and stationary cells. The probability that the PC1 value of a wild-type exponential cell is lower than the PC1 value of a stationary RFP cell is 99.9% (AUC = 0.999), while the probability that the PC1 value of a GFP exponential cell is lower than the PC1 value of a stationary RFP cell is 99.67% (AUC = 0.9967). Thus, for the GFP exponential cells, 23 out of 10,000 cell pairs (1 exponential, 1 stationary) will be incorrectly ranked due to cross-contamination in the GFP cells (likely from ambient cDNA or multiplets). We conclude that cross-contamination has a minor effect on the capacity for PETRI-seq to distinguish between populations. Finally, we confirmed that in the original library for Experiment 1.10 (before down-sampling UMIs and selecting a subset of cells), the relative representation of UMIs from exponential and stationary cells (as predicted by the PCA) were roughly equal (50.3% stationary, 45.6% exponential), indicating that the cross-contamination analysis for the GFP exponential population would be reciprocal for the RFP stationary population.

**Figure S12:**
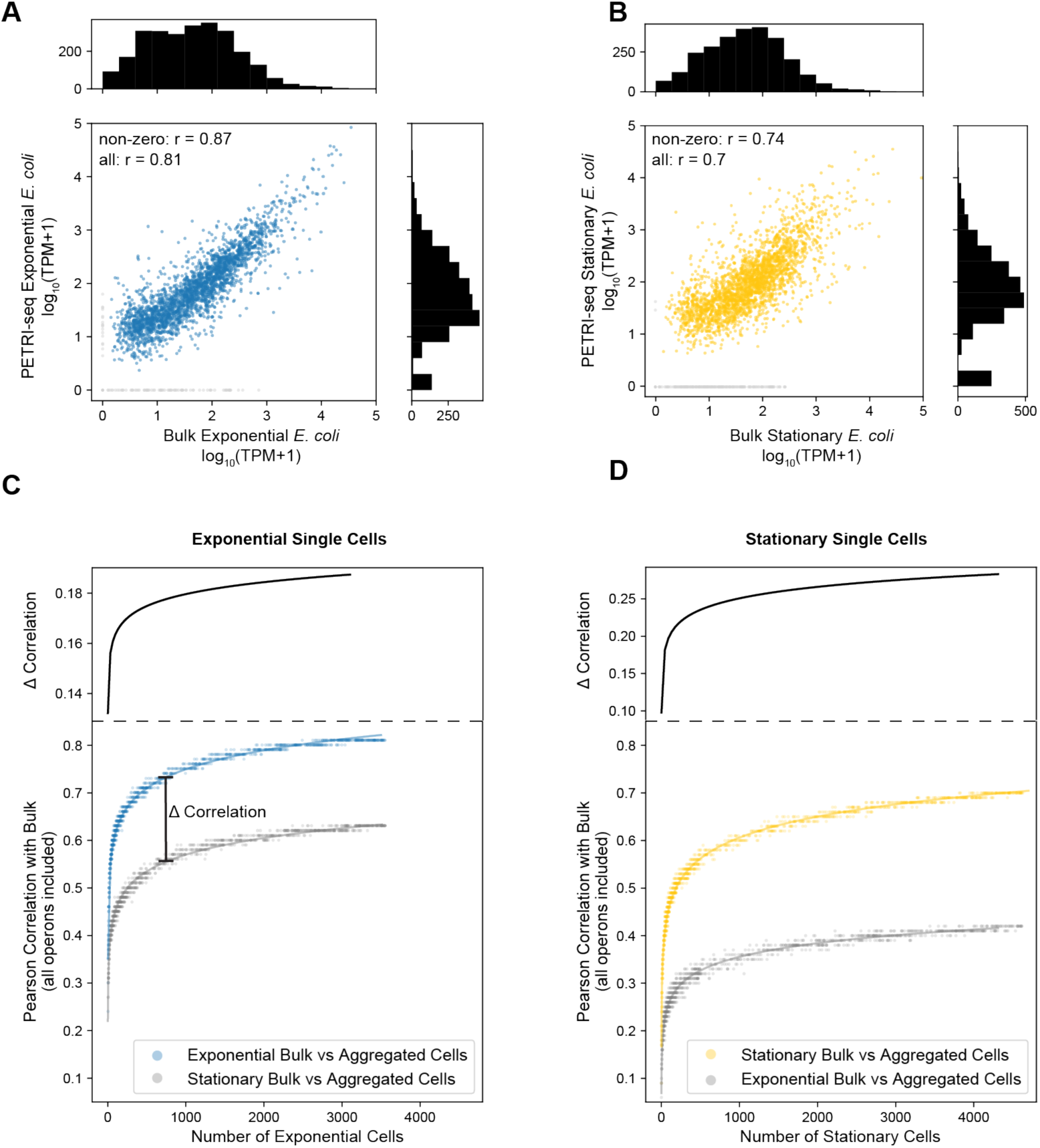
Defining consensus transcriptional states of sub-populations using PETRI-seq. (**A**) Correlation between mRNA abundances from 3,547 aggregated wild-type exponential cells (Experiment 2.01) vs. bulk preparation from fixed exponential GFP-expressing *E. coli* cells. The Pearson correlation coefficient (r) was calculated for 2,458 out of 2,612 total operons, excluding those with zero counts in either library (grey points). If all operons are included, r = 0.81. (**B**) Correlation between RNA abundances from 4,627 aggregated wild-type stationary cells (Experiment 2.01) vs. bulk preparation from fixed RFP-expressing stationary *E. coli* cells. The Pearson correlation coefficient (r) was calculated for 2,362 out of 2,612 total operons, excluding those with zero counts in either library (grey points). If all operons are included, r = 0.7. Bulk libraries in (A) and (B) were prepared from different cultures (on different days) than PETRI-seq libraries. Importantly, stationary populations on different days were often biologically distinct from each other (Figure S11A,B), though they also exhibited similar expression patterns relative to exponential cells. (**C**) *Bottom*: The correlation between the aggregated mRNA counts of single exponential cells (PETRI-seq) and an independently prepared bulk exponential population increases as more single cells are included. Correlations were calculated from log10(TPM+1) for each sample. Single cell transcriptomes were prepared by PETRI-seq, and cell states were predicted by PCA. Best fit line is shown behind original data points (y = ln(x) + b, r > 0.99). *Top*: Difference between the y-values of the best-fit lines for the top curve and bottom curve in plot below. (**D**) *Bottom*: The correlation between the aggregated mRNA counts of single stationary cells (PETRI-seq) and an independently prepared bulk stationary population increases as more single cells are included. Correlations were calculated from log10(TPM+1) for each sample. Single cell transcriptomes were prepared by PETRI-Seq, and cell states were predicted by PCA. Best fit line is shown behind original data points (y = ln(x) + b, r > 0.98). *Top*: Difference between the y-values of the best-fit lines for the top curve and bottom curve in plot below.

**Figure S13:**
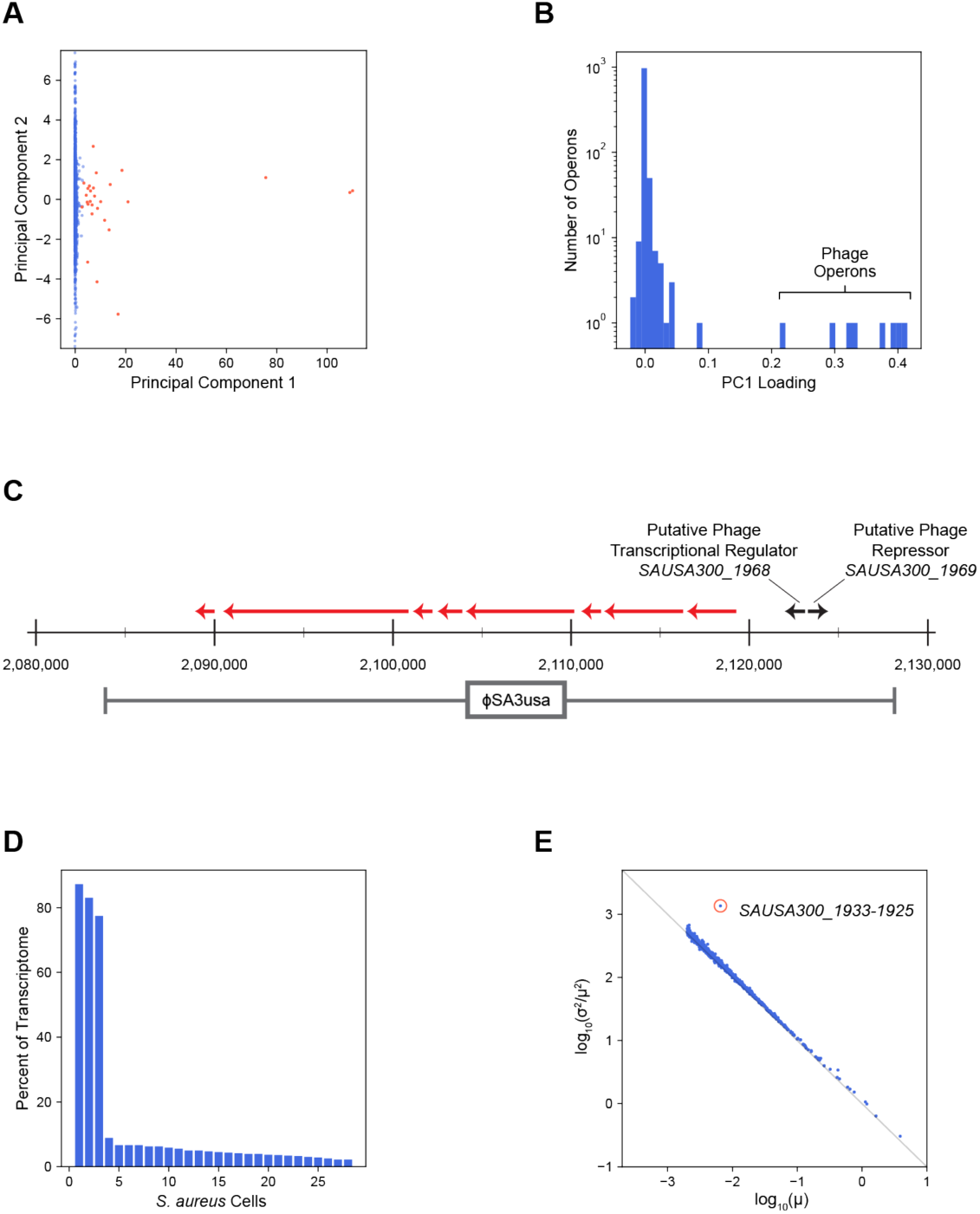
PETRI-seq reveals rare prophage induction in sub-population of *S. aureus* cells. (**A**) *S. aureus* cells plotted on PC1 and PC2. 5,604 cells are included. A small population of 28 cells (*red*) expressed operons from the φSA3usa phage. (**B**) Distribution of PC1 loadings for all operons included in the *S. aureus* analysis. Eight operons from the φSA3usa phage have the highest PC1 loadings. (**C**) Map of genomic region surrounding φSA3usa in the genome of *S. aureus* strain USA300. Red arrows indicate phage operons upregulated along PC1. (**D**) Percent of mRNA UMIs mapped to the φSA3usa phage for the 28 cells containing phage UMIs. Three cells are composed of >77% phage transcripts. (**E**) Noise (σ^(^/μ^(^) versus mean (μ) for operon expression within an *S. aureus* population of 6,663 cells. 676 operons are included, as operons with fewer than 6 raw total UMIs and a mean less than 0.002 after normalization were excluded. The circled operon (red) is *SAUSA300_1933-1925*, which deviated significantly from the rest of the distribution (z-score = 20.6, p < 10^-90^, FDR < 0.01).

**Figure S14:**
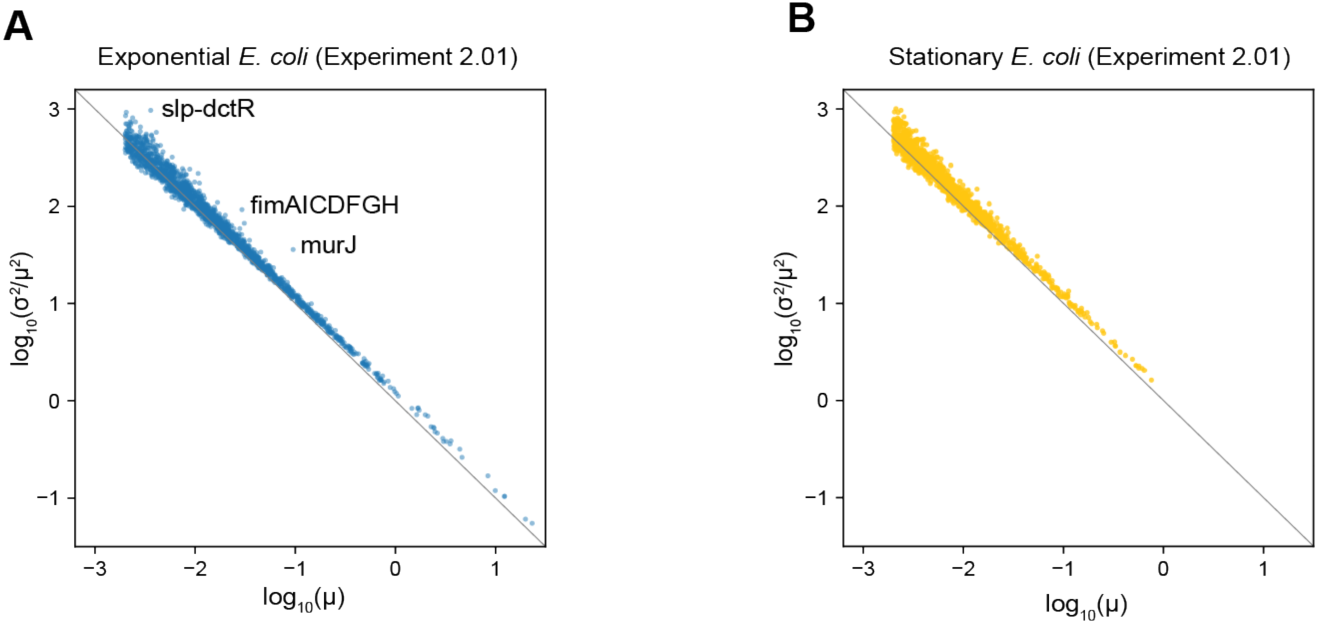
Analysis of hyper-variable operons within *E. coli* populations. (**A,B**) Noise (σ^(^/μ^(^) versus mean (μ) for operon expression in either exponential (A) or stationary (B) cells from Experiment 2.01. Lines at y = -x indicate Poisson noise where σ^2^ = μ. Operon counts were normalized for each cell before plotting. Operons with fewer than 6 raw total UMIs and a mean less than 0.002 after normalization were excluded, resulting in 1,960 operons in (A) and 1,219 operons in (B). Five operons significantly (FDR < 0.01) deviated from the other operons in (A): *sip-dctR* (z-score = 7.3, p = 3*10^-13^), *murJ* (z-score = 6.7, p = 3*10^-11^, *fimAICDFGH* (z-score = 5.4, p = 7*10^-8^), *mdtL* (z-score = 4.8, p = 1*10^-6^), *rnhA* (z-score = 4.6, p = 4*10^-6^). *fimAICDFGH*, which encodes the type I fimbriae system, has been shown previously to exhibit population-level phase variation that is mediated by transcriptional control^39^.

**Table S1:**
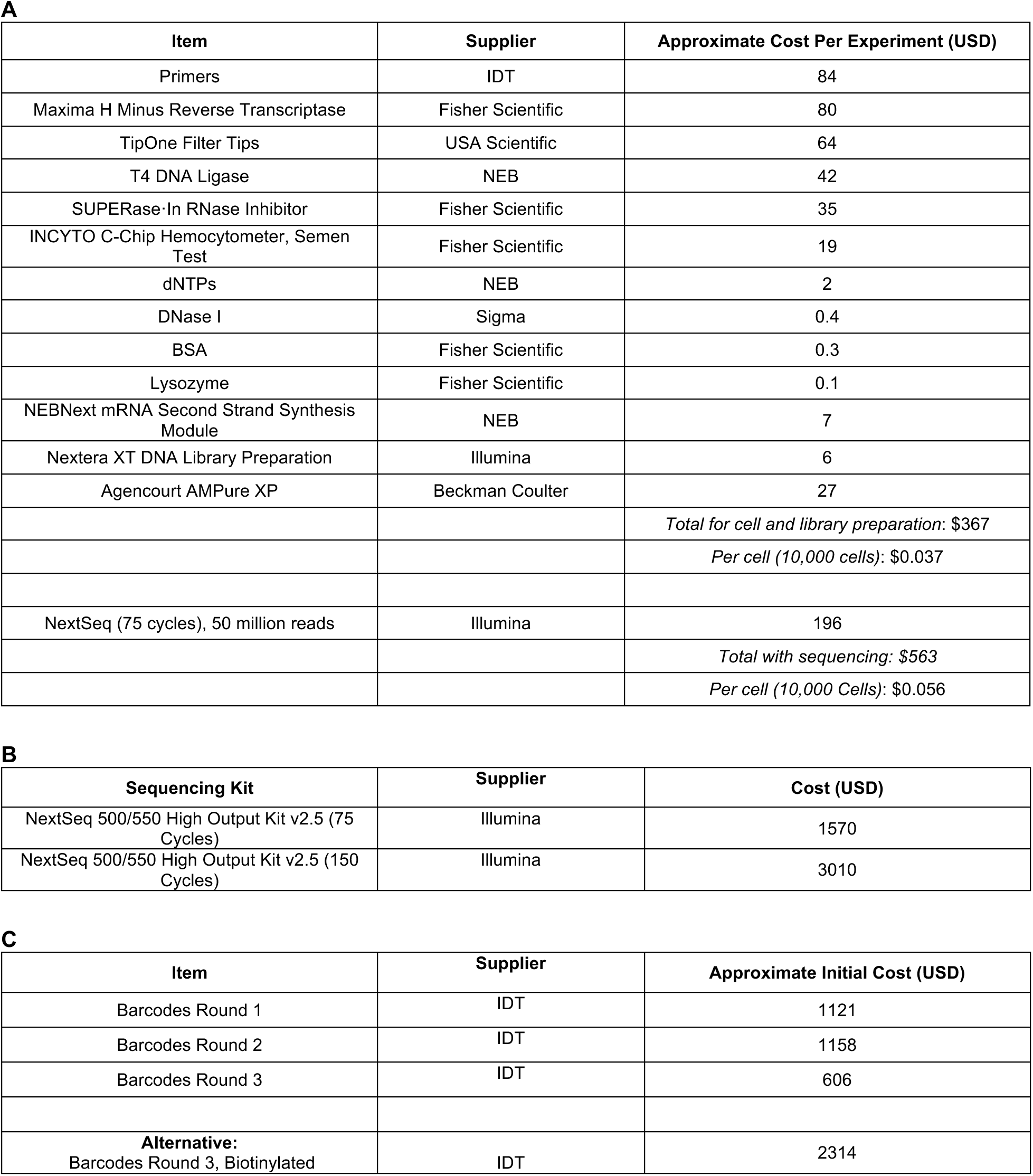
C**o**st **Breakdown for PETRI-Seq**

| Item | Supplier | Approximate Cost Per Experiment (USD) |
| --- | --- | --- |
| Primers | IDT | 84 |
| Maxima H Minus Reverse Transcriptase | Fisher Scientific | 80 |
| TipOne Filter Tips | USA Scientific | 64 |
| T4 DNA Ligase | NEB | 42 |
| SUPERase-In RNase Inhibitor | Fisher Scientific | 35 |
| INCYTO C-Chip Hemocytometer, Semen Test | Fisher Scientific | 19 |
| dNTPs | NEB | 2 |
| DNase I | Sigma | 0.4 |
| BSA | Fisher Scientific | 0.3 |
| Lysozyme | Fisher Scientific | 0.1 |
| NEBNext mRNA Second Strand Synthesis Module | NEB | 7 |
| Nextera XT DNA Library Preparation | Illumina | 6 |
| Agencourt AMPure XP | Beckman Coulter | 27 |
| | | *Total for cell and library preparation: \$367* |
| | | *Per cell (10,000 cells): \$0.037* |
| NextSeq (75 cycles), 50 million reads | Illumina | 196 |
| | | *Total with sequencing: \$563* |
| | | *Per cell (10,000 Cells): \$0.056* |

**B**
| Sequencing Kit | Supplier | Cost (USD) |
| --- | --- | --- |
| NextSeq 500/550 High Output Kit v2.5 (75 Cycles) | Illumina | 1570 |
| NextSeq 500/550 High Output Kit v2.5 (150 Cycles) | Illumina | 3010 |

**Table S1: Cost Breakdown for PETRI-Seq**
| Item | Supplier | Approximate Initial Cost (USD) |
| --- | --- | --- |
| Barcodes Round 1 | IDT | 1121 |
| Barcodes Round 2 | IDT | 1158 |
| Barcodes Round 3 | IDT | 606 |
| **Alternative:**<br>Barcodes Round 3, Biotinylated | IDT | 2314 |

**Table S2:**
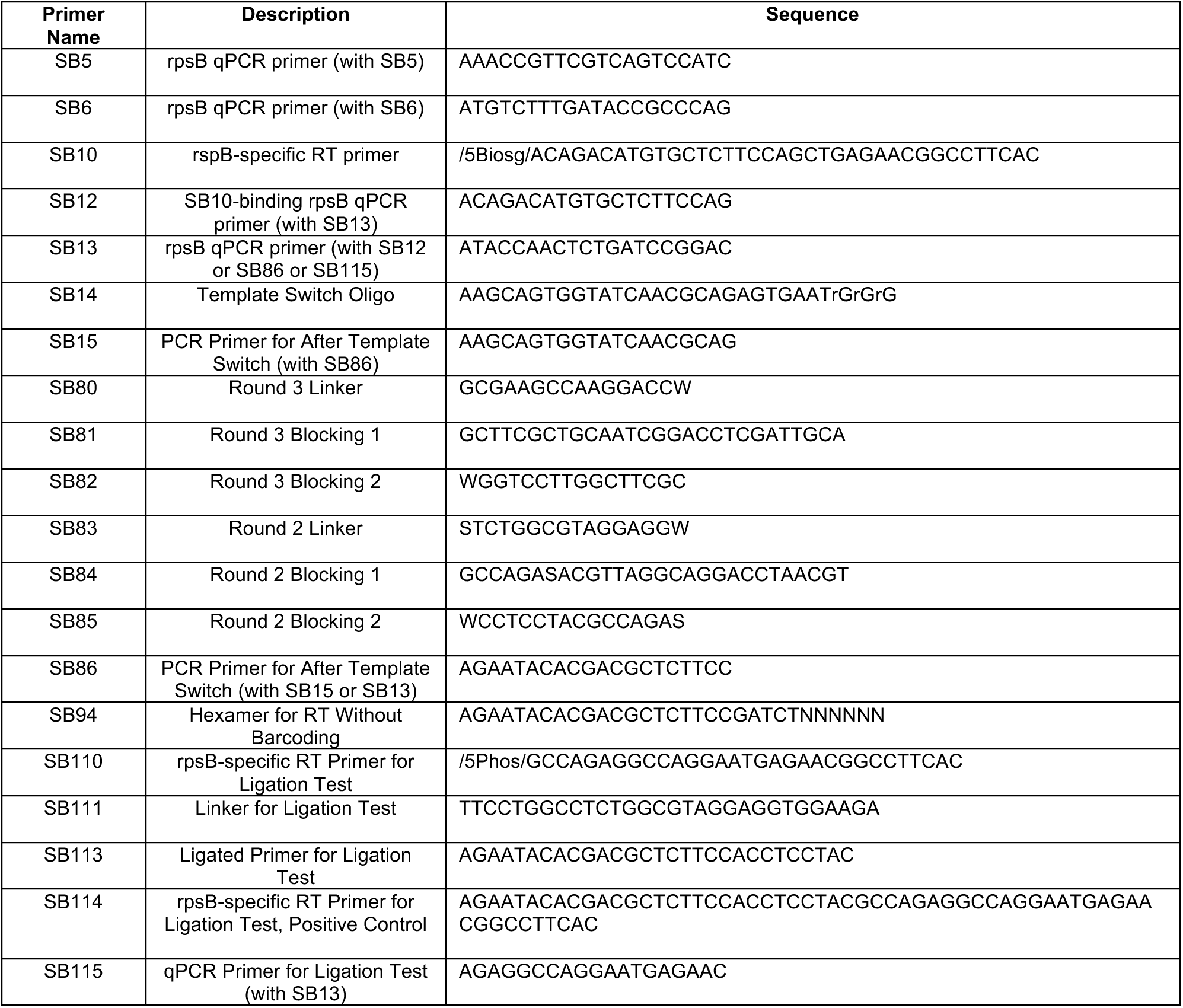
O**l**igonucleotides **used in this study** Name, description, and sequence of all single-tube (excluding 96-well barcode plates) oligonucleotides used in the study.

## Additional Files

**Table S3: 96-Well oligonucleotides used for PETRI-seq barcoding** Sequences of 96 round 1 RT primers, 96 round 2 ligation primers, and 96 round 3 ligation primers.

**Table S4: Overview of experiments included in this study** Table includes cell types, thresholds, number of cells, figures used in, total number of reads, percent of reads aligned, and mean reads per cell.

**Table S5: Supplementary statistical data for Figure S3 and Figure S8**

**Table S6: Count matrix for Experiments 1.06SaEc, 1.10, 2.01 and Bulk Libraries** BC by operon matrix of raw UMI counts for the three primary experiments used in this study. Anti-sense operons were excluded. BCs with prefix “SB346” are from 1.06SaEc, “394A” from 1.10, and “SB442” from 2.01. Bulk libraries for stationary RFP-expressing *E. coli* cells (“SB369”) and exponential GFP-expressing *E. coli* cells (“SB371”) are also included; reads, rather than UMIs, are reported for bulk libraries. Operon names with prefix “U00096:” originate from *E. coli*, while operons with prefix “CP000255:” originate from *S. aureus*.

